# Fusion of the N-terminal 119 amino acids with the RelA-CTD renders its growth inhibitory effects ppGpp-dependent

**DOI:** 10.1101/2021.03.21.436043

**Authors:** Krishma Tailor, Prarthi Sagar, Keyur Dave, Jayashree Pohnerkar

**Affiliations:** Department of Biochemistry, Faculty of Science, Maharaja Sayajirao University of Baroda, Vadodara – 390002, Gujarat, India; Department of Biochemistry and Molecular Biology, Howard University College of Medicine, Washington, DC, U.S.A

**Keywords:** (p)ppGpp, RelA, HD domain, RelA-CTD

## Abstract

The guanosine nucleotide derivatives ppGpp and pppGpp, are central to the remarkable capacity of bacteria to adapt to fluctuating environment and metabolic perturbations. These alarmones are synthesized by two proteins, RelA and SpoT in *E. coli* and the activities of each of the two enzymes are highly regulated for homeostatic control of (p)ppGpp levels in the cell. Although the domain structure and function of RelA are well defined, the findings of this study unfold the regulatory aspect of RelA that is possibly relevant *in vivo*. We uncover here the importance of the N-terminal 1-119 amino acids of the enzymatically compromised (p)ppGpp hydrolytic domain (HD) of monofunctional RelA for the (p)ppGpp mediated regulation of RelA-CTD function. We find that even moderate level expression of RelA appreciably reduces growth when the basal levels of (p)ppGpp in the cells are higher than in the wild type, an effect independent of its ability to synthesize (p)ppGpp. This is evidenced by the growth inhibitory effects of oversynthesis of the RelA-CTD in the *relA*^+^ strain but not in *relA* null mutant, suggesting the requirement of the functional RelA protein for basal level synthesis of (p)ppGpp, accordingly corroborated by the restoration of the growth inhibitory effects of the RelA-CTD expression in the *relA1 spoT202* mutant. The N-terminal 119 amino acids of RelA fused in-frame with the RelA-CTD, both from 406-744 amino acids (including TGS) and from 454-744 amino acids (sans TGS) caused growth inhibition only in *spoT1* and *spoT202 relA1* mutants, uncovering the hitherto unrealized (p)ppGpp-dependent regulation of RelA-CTD function. An incremental rise in the (p)ppGpp levels is proposed to progressively modulate the interaction of RelA-CTD with the ribosomes, with possible implications in the feedback regulation of the N-terminal (p)ppGpp synthesis function, a proposal that best explains the nonlinear relationship between (p)ppGpp synthesis and increased ratio of RelA:ribosomes, both *in vitro* as well as *in vivo*.

## Introduction

Environmental stressors elicit largely conserved adaptive responses in bacteria (and plants), mediated and coordinated by the hyperphosphorylated nucleotides, ppGpp and pppGpp (together called (p)ppGpp). These signalling nucleotides control various cellular activities at transcriptional, translational, and posttranslational levels (Hauryliuk *et al*., 2015). Recent studies have realized that the intracellular pool size of (p)ppGpp does not act as a biphasic switch rather, incremental levels of (p)ppGpp exert differential effects on cell physiology including its role in virulence, pathogenesis, antibiotic resistance/tolerance, sporulation, biofilm and persisters’ cell formation (Dozot *et al*., 2006, Geiger *et al*., 2010, Ochi *et al*., 1981, Poole, 2012, Schofield *et al*., 2018); this is besides the well characterised stringent response (Cashel *et al*., 1996, Potrykus & Cashel, 2008).

These nucleotides derivatives are metabolised by two types of proteins – widely distributed, highly conserved, long multidomain, bifunctional RSH (RelA SpoT Homolog) and monofunctional small alarmone synthetases (SASs) and hudrolase (SAHs) which are limited in their distribution to Gram positive Firmicutes and *Actinobacteria* (Atkinson *et al*., 2011, Gaca *et al*., 2015, Ronneau & Hallez, 2019, Steinchen & Bange, 2016). In *E*. *coli* and in other γ-proteobacteria, (p)ppGpp is synthesised by long monofunctional RSH, RelA, whose hydrolytic domain is heavily compromised; whereas, the second RSH protein, SpoT is a bifunctional enzyme containing both synthetase and hydrolase activities and normally function in homeostatic regulation of (p)ppGpp levels by hydrolysing excess (p)ppGpp (An *et al*., 1979, Hernandez & Bremer, 1991). SpoT is nevertheless a weak (p)ppGpp synthetase, responsive to signals of fatty acid, iron and carbon limitation (Iyer *et al*., 2018, Seyfzadeh *et al*., 1993, Spira *et al*., 1995, Vinella *et al*., 2005, Wang *et al*., 2016, Xiao *et al*., 1991); interestingly though, fatty acid starvation induced rapid synthesis of (p)ppGpp has recently been shown to lead in part to amino acid starvation and contribution from RelA followed by its delayed synthesis by SpoT (Gentry & Cashel, 1996, Sinha *et al*., 2019). The characteristic functional elements of the long RSH enzyme (RelA/SpoT and Rel) includes a N-terminal enzymatic domain harbouring motif for hydrolase and/or (p)ppGpp synthetase activity and a C-terminal regulatory domain (CTD) containing conserved motifs - TGS (ThrRS, GTPases and SpoT), DC (aspartate-cysteine motif) and ACT (aspartate kinase, chorismate mutase and TyrA) also called RRM (RNA recognition motif) (Atkinson *et al*., 2011). Basic regulation of (p)ppGpp synthesis function of long RSH enzymes is quite well conserved across mono- and bifunctional long RSH proteins which is reproduced *in vitro* (Arenz *et al*., 2016, Brown *et al*., 2016, Haseltine & Block, 1973) (Kushwaha *et al*., 2019, Loveland *et al*., 2016, Winther *et al*., 2018). The CTD of the long RSH proteins has a unique regulatory role of inhibiting the (p)ppGpp synthetase function of the NTD but not the hydrolytic activity by the conserved aspartate-cysteine motif (DC) that has been also proposed to be involved in the oligomerization of RelA in the ribosome unbound state (Gratani *et al*., 2018, Yang & Ishiguro, 2001, Avarbock *et al*., 2005, Butland *et al*., 2008, Gropp *et al*., 2001); the activation of ribosome-bound RelA is proposed to be due to the association of monomeric RelA with the ribosome and deacylated tRNA which relieves the autoinhibitory effect of the CTD on the NTD (Arenz *et al*., 2016) (Avarbock *et al*., 2005) (Jain *et al*., 2006b, Loveland *et al*., 2016). Furthermore, recent studies indicated an allosteric regulation of RelA, both positive and negative by the end product, (p)ppGpp. The regulatory effectors of RelA enzymatic activity for (p)ppGpp synthesis include positive end product activation by pppGpp at low (∼100uM), and negative inhibitory effect at higher (∼400 uM) concentration (Kudrin *et al*., 2018, Wendrich *et al*., 2002, Shyp *et al*., 2012); the site for positive activation has been localized to the N-terminal segment (NTD) segment of Rel of *B. subtilis*, *S*. *equisimilis, M. tuberculosis*, *M. smegmatis* and *Francisella tularensis* (Jain *et al*., 2006a, Wilkinson *et al*., 2015, Takada *et al*., 2021) and RelA of *Escherichia coli* (Kudrin *et al*., 2018, Shyp *et al*., 2012, Takada *et al*, 2021). Interestingly, (p)ppGpp-mediated regulation of synthase activity of Rel by the Rel-CTD in *M. smegmatis* has also been demonstrated. The mechanism of negative feedback regulation has been proposed to be important for fine tuning the Rel’s (p)ppGpp synthesis function (Syal *et al*., 2015).

The results presented here provide evidence for the existence of regulation by (p)ppGpp of the RelA-CTD function in *E. coli*. The novel aspect of regulation of RelA’s growth inhibitory function became apparent with the fortuitous cloning of RelA-CTD sequence remaining after the removal of the DNA between two *Pvu*II sites at positions 354 and 1362 in the *relA* gene sequence which also results in an in-frame fusion of the N-terminal 1- 119 amino acids to 454-744 amino acid sequence of the RelA-CTD. We showed that the low to moderate level expression of RelA-CTD fusion gene significantly inhibits growth, albeit only when the basal levels of (p)ppGpp are higher than in the wild type. Thus, the combination of mutations, *relA*^+^ *spoT1* and *relA1 spoT202* which enhances intracellular levels of (p)ppGpp more than that by *relA1 spoT1* or *relA*^+^ *spoT*^+^, was effective in promoting inhibition of the growth by fusion RelA-CTD, substantiating the growth inhibitory effects to be dictated by the intracellular levels of (p)ppGpp and independent of RelA’s (p)ppGpp synthesis ability. This result was subsequently confirmed by PCR based in-frame fusion of 1-357 nucleotides (119 amino acids) to 1014-2235 nucleotides of RelA-CTD (406-744 amino acids) which additionally contains TGS domain. Our results are compatible with the proposal that (p)ppGpp mediates, through the N-terminal 119 amino acids, regulation of binding of RelA-CTD to the target ribosomes, presumably important in the feedback regulation of RelA’s (p)ppGpp synthesis function. Recently, very high-level expression of RelA-CTD has been shown to inhibit the growth, irrespective of the *relA* genotype of the host as a result of the decrease in the rate of protein synthesis and interference in the normal stringent response (Gropp *et al*., 2001), whereas moderate levels are without an effect on growth (Turnbull *et al*., 2019). The significance of our results is discussed.

## Results

The genesis of the present work was an attempt to test the functional significance of the synteny of *relA* (encoding (p)ppGpp synthetase I) and *rumA* (coding for 23S rRNA methyl transferase) in genomes of several gram-negative bacteria, including some important pathogens. Incidentally, the U1939 of 23S rRNA is placed close to the acceptor arm of the uncharged tRNA where RelA binds on the ribosome (Yusupov *et al*., 2001, Persaud *et al*., 2010). Persaud *et.al*., (2010) addressed the functionality of *rumA* and reported *rumA* to be inessential for growth as the null mutation in the *rumA* gene is not associated with any growth defect. Since their mutation in *rumA* was a deletion of its ORF, and since all the promoters of *relA* are present in the upstream *rumA* gene (Fig.1), both transcriptional and translational regulation of *relA,* if there were any, are expected to be abolished in the mutant. We tested the essentiality of *rumA* in this work by instead creating an insertion of chloramphenicol acetyl transferase (*CAT*) or Kanamycin resistance (*KAN*) cassette in the *rumA* ORF at a unique *Sal*I site −878 bp upstream of all the known promoters of *relA* gene (Fig.1). Interestingly, we find that the *CAT* insertion in one of the two orientations exerted a *cis* effect on expression of *relA* gene, causing elevated synthesis of RelA protein, whereas, insertion of the *CAT* gene in the opposite orientation, and the insertion of *KAN* gene cassette in either orientation, did not exert any effect on expression of *relA* (see below), validating the result that lack of methylation at U1939 of 23SrRNA by RumA is not relevant for *relA* regulation (Persaud *et al*., 2010). We investigated in this paper the *cis* effect of the *CAT* insertion mutation that increases *relA* expression and its phenotypic consequences.

**Fig. 1:**
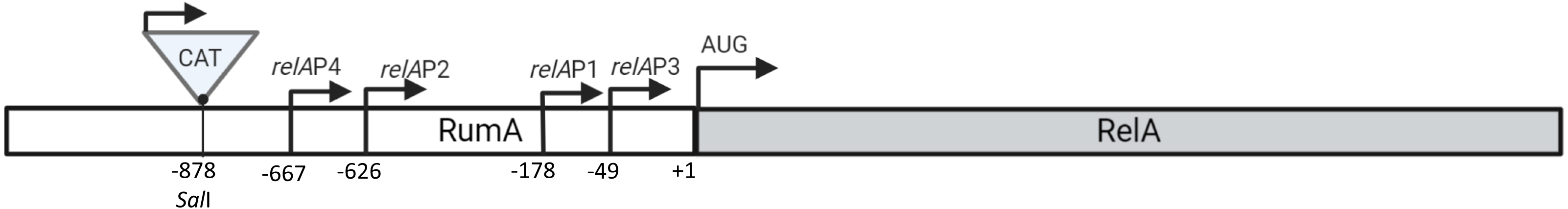
Genetic organization and transcriptional signals of the *relA* gene in the upstream *rumA* gene on *E coli* chromosome. The four arrows (P1, P2, P3, and P4) indicate the known *relA* promoters and the numbers represents the +1 start position of the transcript from each of the four promoter in the *rumA* gene. The triangle indicates *CAT* antibiotic cassette insertion at the *Sal*I site −878 bp upstream of all the known promoters of the *relA* gene. The arrow on *CAT* cassette indicates transcription direction of *CAT*. The AUG represents start site of *relA* translation.

Because of the intrinsic variation in the basal levels of (p)ppGpp in several *Escherichia coli* strains due to polymorphism at *relA* and *spoT* (Laffler & Gallant, 1974a, Brown *et al*., 2002, King *et al*., 2004, Spira & Ferenci, 2008, Ferenci *et al*., 2011), we used two representative strains of *E. coli*, namely MG1655 and MC4100 which are recognized as being (p)ppGpp sensitive and tolerant respectively, in terms of (p)ppGpp related phenotypes being accentuated in the latter strain. We performed most of the experiments with both the strains and found a large part of the results to be same between them. To avoid repetition, the results described here are reported for the strain MG1655 unless indicated otherwise. Briefly, genotypes relevant to (p)ppGpp homeostasis for each of the strain are: MG1655 is wildtype at both *relA* and *spoT* loci, whereas MC4100 has two mutations, *relA1* and *spoT1*. The *relA1* mutation is an *IS2* insertion between 85^th^ and 86^th^ codon of *relA* gene, as a result, the mutant protein retains 1% residual RelA activity (Metzger *et al*., 1989). The *spoT1* allele contains a substitution (H255Y) in the synthesis domain and a two-amino acid insertion between residues 82 and 83(+QD) in the hydrolysis domain of the SpoT protein (Spira *et al*., 2008). The compromised (p)ppGpp synthetic and hydrolytic functions of SpoT1 are responsible for slow (20-fold) decay in the first order kinetics of (p)ppGpp during stringent response as well as a severe impairment in (p)ppGpp degradation during exponential growth, resulting in its higher basal levels (Laffler & Gallant, 1974a, Laffler & Gallant, 1974b, Fiil *et al*., 1977, Sarubbi *et al*., 1988). However, the reasons for sensitivity and tolerance of strains MG1655 and MC4100 respectively to (p)ppGpp mediated effects are more than the differences at *relA* and *spoT* (Spira & Ferenci, 2008).

### Insertion of *CAT*/*KAN* cassette in *rumA* gene

*rumA* mutants were generated by the strategy of recombineering (Datsenko & Wanner, 2000) of *CAT/KAN* cassette in *rumA* DNA. (i) *CAT* cassette was inserted at the unique *SalI* restriction enzyme site −878 bp upstream of *relA* sequence in the orientation where the direction of transcription/translation of *CAT* is same as that of *relA* in the MG1655/MC4100 *relA*^+^ background. This insertion is upstream of all the known promoters of *relA* mapped in *rumA* gene sequence (Fig.1), and the only one to affect the growth phenotype of the mutant (see below). (ii) The *CAT* insertion at *SalI* restriction enzyme site in the opposite orientation in KP59 and KP60 (Fig.3) or (iii) in either orientation at the unique *MluI* restriction enzyme site (−1310 bp) in *rumA* DNA (KP9, KP25) did not have any effect on growth (and *relA* synthesis) (Fig.3). (iv) We next tested if *CAT* gene replacement by *KAN* gene cassette affects growth if its direction of transcription/translation were same as that of the *relA*. The *KAN*^r^ cassette was cloned at *SalI* site in either orientation in the *rumA* DNA by the recombineering method same as that for the *CAT* gene (see Materials and Methods). However, the *KAN*^r^ insertion was without any effect on the growth/expression of RelA (data not shown). The phenotypic effects of orientation specific insertion of *CAT* gene in *rumA* can be due to the chance generation of a promoter at the unique junction between the *CAT* and *rumA* sequences. We did not study this proposal further.

We describe below the phenotypes of the effect of increased synthesis of RelA in KP32 (*spoT1 rumASalI*::*CAT*). The insertion mutation in *rumA* is hereafter referred to as *rumA*::*CAT*.

### The requirement of *spoT1* mutation is obligatory for manifestation of the *rumA*::*CAT* phenotype

The fact that *spoT1* mutation is required for the phenotype became evident when the effect of the mutation was manifest in KP4 (MC4100 *relA*^+^ *spoT1*) but not in MG1655 strain. We constructed *spoT1* derivative of MG1655 by cotransduction with linked Δ*pyrE748*::*KAN*. P1 lysate prepared on KP11 (MC4100 *spoT1*Δ*pyrE748*::*KAN*) was used to transduce *spoT1* mutation in MG1655 strain. The *spoT1* mutation was verified by DNA sequencing (data not shown). In contrast to KP4 (MC4100 *relA*^+^ *spoT1*) strain, MG1655 *spoT1* transductant (KP31) was marginally affected for growth in minimal medium (Fig.2A) (Sarubbi *et al*., 1988).

**Fig. 2:**
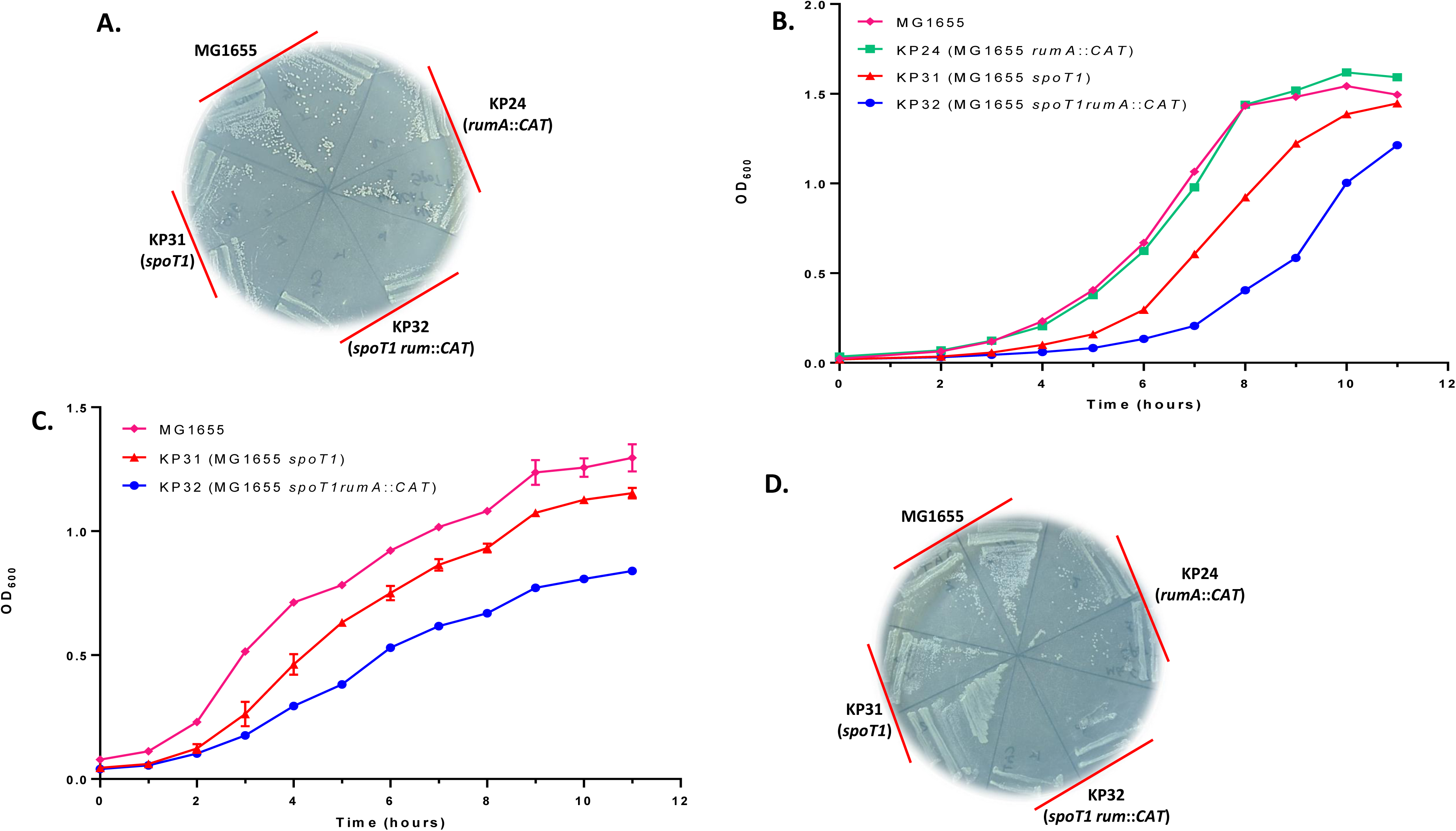
*rumA*::*CAT* mutant grows poorly on both minimal medium and on amino acid starvation plates: (**A**) Growth of the *rumA* mutant on minimal agar plate at 30°C. (**B**) Growth curve measurement of *spoT1 rumA* and *spoT1* mutants of MG1655 strains in minimal broth at 37°C and in (**C**) LB at 30°C. Growth curve is representative of three independent experiments. (**D**) The growth of *rumA* mutant on minimal agar plate supplemented with 3-AT.

### Phenotypes of overexpression of *relA*

#### (i) The *rumA*::*CAT* mutant (KP32) grows unusually slowly on M9 minimal medium

Of the two important phenotypes of the *rumA*::*CAT* mutant described here, the first pertains to the growth of the mutant on M9 minimal agar plate. The slow growth of KP32 (*spoT1 rumA*::*CAT*) is striking in that there is no colony formation for ∼24 hours of incubation on minimal medium at 37°C (Fig.2A). This slow growth phenotype was confirmed by measuring steady state growth rates of different mutants (Fig.2B). Following nutrient shift down from LB to M9 broth, KP32 (*spoT1 rumA*::*CAT*) mutant reproducibly exhibited a long lag of approximately 7-9 hours before the cells’ doubling could be recorded. The *spoT1* derivative of MG1655, KP31, took 2-3 hours under the same condition (data not shown). The growth in the nutrient rich LB medium was slow as well, more so at 30°C than at 37°C (Fig.2C). The markedly slow growth of KP32 (*spoT1 rumA*::*CAT*) and KP8 (MC4100 *relA*^+^ *spoT1 rumA*::*CAT*) mutant often yielded fast growing suppressors (see below).

### The *rumA*::*CAT* mutant (KP32) is unable to express normal stringent response

KP32 (*spoT1 rumA*::*CAT)* grows poorly on minimal agar plate supplemented with 3-AT. Unlike the wild type *relA*^+^ strain, the *rumA*::*CAT* mutant, though genetically *relA*^+^, grows one notch better than the *relA* null mutant control on amino acid starvation plate (Fig.2D), the growth is same as on minimal medium as if ‘frozen’. We confirmed by DNA sequencing that the *relA* gene in KP32 does not contain any mutation; also replacing the *spoT1* mutation with *spoT*^+^ by P1 transduction abrogates the unusual stringent response of the KP32 mutant and the *spoT*^+^ *rumA*::*CAT* transductant (KP24) behaves normally as *relA*^+^ strain with respect to growth on minimal plate, as indicated earlier (Fig.2A & B) and on minimal plate supplemented with 3-AT (Fig.2D). We present below the evidence that RelA is overexpressed in the *rumA* insertion mutant, KP32 which explains most of its phenotypes.

### Isolation of spontaneous suppressors of the slow growth phenotype of *rumA*::*CAT* mutant

Growth of *rumA*::*CAT* mutant was severely affected on minimal medium to the extent that this growth defect could be used in selection of suppressor mutants. Fast growing colonies were often seen on the plate within 24 hours of incubation. The suppressors of KP8 (MC4100 *relA*^+^ *spoT1 rumA*::*CAT*) were invariably *relA* null mutants as they did not grow on amino acids starvation plates containing 3-AT (Fig.3B), and unlike KP4 (MC4100 *relA*^+^) or KP8 (MC4100 *relA*^+^ *spoT1 rumA*::*CAT*), grew as fast as the parent MC4100KP (*relA1*) on minimal medium (Fig.3A). Further, we transduced *relA1* mutation from MC4100KP to KP8 (MC4100 *relA*^+^ *spoT1 rumA*::*CAT*) in P1 transduction using the linked *cysI*::*Tn*10*kan*. One out of 10,000 *Kan*^r^ transductants was *relA1*, exhibiting the characteristic fast growth on minimal medium (Fig.3A) and unable to grow on starvation plate (3-AT) (Fig.3B). Presence of *relA1* (*relA*::*IS*2) and *rumA*::*CAT* mutations was confirmed by PCR (data not shown).

**Fig. 3:**
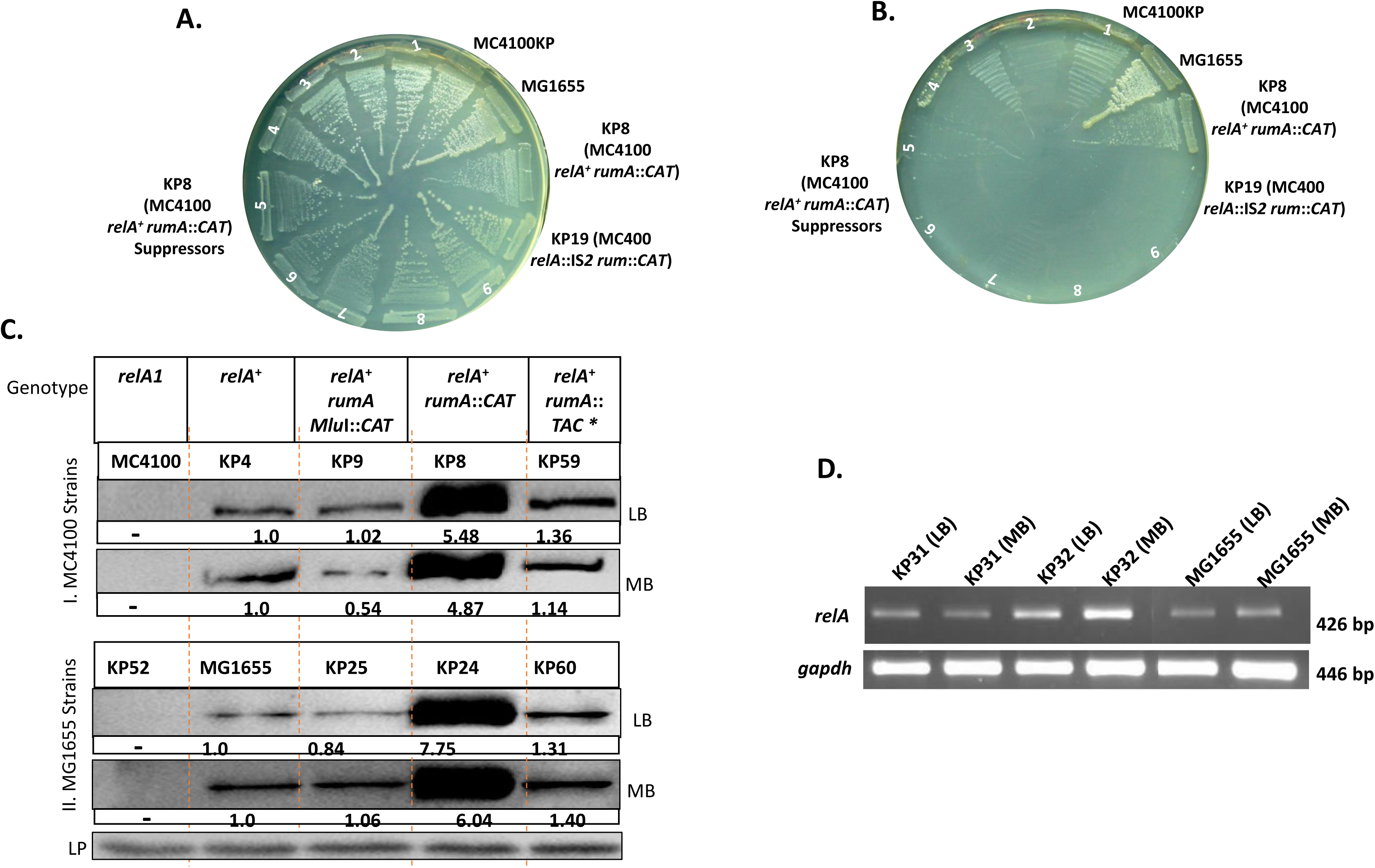
Moderate level expression of RelA in the context of *spoT1* mutation is the cause of phenotypes of *rumA*::*CAT* mutant. Suppressors of slow growth of the *rumA*::*CAT* mutant are RelA**^-^**. The growth phenotype of fast-growing variants of KP8 was assessed on (**A**) minimal agar plate and (**B**) minimal agar plate supplemented with 3-AT. The plates were examined after 24 hours of incubation at 37°C. **(C) Quantification of RelA protein by Western blot.** RelA protein was measured in cell extract of **(I)** *rumA* mutant derivatives of MC4100 grown in rich media (LB) and in minimal media (MB); **(II)** cell extracts of *rumA* mutant derivatives of MG1655 grown in LB and in MB at 37°C. Upper panel represents genotypes of mutants of MC4100 **(I)** and MG1655 **(II)** derivatives. Fold differences in protein levels of *rumA*::*CAT* relative to wild type represents average of the three independent experiment and indicated against each lane. In immunoblots of MG1655 strains, the lower panel (LP) shows β-Galactosidase protein as an internal protein control detected by anti-β-gal antibody. (*) Insertion of the CAT cassette in the *rumA* gene (*rumA::TAC*) in the orientation opposite to that of *rumA::CAT*. **(D) Semi-quantitative measurement of *relA* transcript by Reverse Transcriptase-PCR.** The cells were harvested for isolation of RNA after their growth in rich media (LB) and minimal media (MB). Upper panel shows 426 bp *relA* amplicon generated using RelA RT primers (Table 2) corresponding to C-terminal portion. Lower panel represents RT-PCR of internal control of *gapdh* mRNA. 100 bp ladder was used as M.W marker.

Surprisingly, the fast growing variants of KP32 (MG1655 *rumA*::*CAT spoT1*) could not be tested for their *relA* status, as *relA1 spoT1* mutant of MG1655 (KP53) was resistant to 3-AT, whereas MG1655 *relA1* mutant (KP52) was sensitive (Fig.S2A). The residual activity of the RelA1 mutant protein is ruled out to be reason of 3-AT resistance of the KP53 mutant because the Δ*PvuII* deletion of the chromosomal *relA* gene corresponding to the N-terminal (p)ppGpp synthetic portion (119-455 amino acids) tagged to the insertion of the *CAT* cassette in the *relA* gene at the unique *SalI* site at position 1619 in the mutant KP54 yielded the same phenotype as *relA1 spoT1* mutant (Fig.S2A). This phenotype is reminiscent of suppression of the sensitivity of *relA* strains of *Salmonella typhimurium* to 3-AT by *spoTI* mutation (Tedin & Norel, 2001). Although the fast-growing mutants of KP31 (MG1655 *spoT1)* could not be directly tested for the RelA^-^ phenotype on 3-AT agar plate, pTE18 *spoT*^+^ transformants of two of the fast-growing mutants were indeed RelA^-^ (sensitive to 3-AT, data not shown).

The result that the slow growth of KP8 (MC4100 *relA*^+^ *spoT1 rumA*::*CAT*) mutant is suppressed by inactivating mutation in *relA* suggests that the growth defect of the strain is presumably due to oversynthesis of RelA. We surmised that the level of RelA protein is increased in the *rumA::CAT* which we confirmed in the experiment described below.

### Measurement of RelA protein by Western blot analysis

We measured RelA protein amounts in different strains grown in both nutrient rich (LB) and minimal medium supplemented with glucose by Western Blot using anti-RelA antibody. From the results of densitometric analysis of more than three independent blots, it is evident that amount of RelA protein is 7-10 fold more in *rumA* mutant than in other strains (Fig.3C). Importantly, we found RelA protein to be increased to similar extent in *rumA*::*CAT* mutant of both MC4100 (*relA*^+^) and MG1655 background under all conditions of growth (Fig.3C). Notwithstanding this increased amounts, as indicated earlier, the reason for the lack of growth phenotype in KP24 (MG1655 *rumA*::*CAT*) is the presence of wild type *spoT*^+^ gene. The increase in the RelA protein in the *rumA*::*CAT* mutant and not in *rumA::TAC* strain (KP59, KP60) is possibly due to increased transcription from a promoter-like sequence at the junction between *rumA* and *CAT* cassette sequence. We therefore estimated *relA* mRNA levels by semiquantitative RT-PCR.

### Amount of *relA* transcript is enhanced in the *rumA*::*CAT* mutant

The *relA* mRNA levels were elevated by ∼10 fold when quantitated by comparative transcriptome analysis of MG1655, KP31 (*spoT1*) and KP32 (*spoT1 rumA*::*CAT*) strains (RPKM - 5:8:67 respectively) and validated by semiquantitative Reverse Transcriptase PCR in the strains grown in LB and in MB. We noted that (i) the *relA* transcript abundance is more in cells grown in MB than in LB. (ii) the transcript level was increased in *rumA*::*CAT* mutant (KP32, KP24) when compared to that in other strains (Fig.3D). Thus, we believe that the insertion of *CAT* cassette in *rumA* in *cis* increases the transcription of the *relA* gene.

### Cloned *spoT*^+^ gene as a multicopy suppressor of the growth defect of KP32 (*spoT1 rumA*::*CAT*)

Genomic library of *E. coli* MG1655 was prepared in pBR322 vector and transformed into KP32 (MG1655 *spoT1 rumA*::*CAT*) mutant. From this library we selected 3 larger size colonies. The DNA prepared from each could complement the slow growth defect of *rumA*::*CAT* mutant, KP8 (MC4100 *relA*^+^ *spoT1 rumA*::*CAT*) and KP32 (MG1655 *spoT1 rumA*::*CAT*) (Fig. 2D) and also returned the normal stringent response of KP8 (Fig.S1A &SB). Nucleotide sequencing revealed that the DNA in each clone contained *spoT*^+^ gene. Subcloned *spoT*^+^ gene was sufficient to complement the growth defect of *rumA* mutant (data not shown).

### Measurement of basal- and amino acid starvation induced (p)ppGpp levels in the *rumA* mutant

Intracellular levels of (p)ppGpp were measured in the cells of MG1655, KP31 (*spoT1*) and KP32 (*spoT1 rumA*::*CAT*) grown in the MOPS minimal medium and also with valine supplementation to cause leucine/isoleucine starvation by the method as described in (Cashel, 1994, Fernández-Coll & Cashel, 2019). The result that there was no demonstrable difference in the (p)ppGpp levels between KP31 (*spoT1*) and KP32 (*spoT1 rumA*::*CAT*) strains was highly reproducible and consistent (Fig.4A, B). This is surprising and unexpected in the context that the *relA* gene cloned downstream of *lacUV5* (*trc*) promoter in the plasmid pALS10/pSM10 synthesizes (p)ppGpp in response to induction of its expression by IPTG (Schreiber *et al*., 1991, Svitil *et al*., 1993) in the absence of starvation for any amino acid. The increase in the (p)ppGpp levels inhibits growth with an approximate linear relationship. The slow growth of the KP32/KP8 (*spoT1 rumA*::*CAT)* mutant was similarly expected to be due to the overexpression of *relA* causing elevated synthesis of (p)ppGpp, compounded by a further enhancement of the intracellular amounts due to its reduced turnover by hydrolysis-defective *spoT1* mutation. Also, there is very little difference, if any, in the levels of (p)ppGpp in different mutants in response to starvation (Fig.4C). The result is important in indicating that there is no defect in the amino acid starvation response in terms of the amount of (p)ppGpp synthesized under starvation condition. Thus, the inadequate growth of the KP32 (*spoT1 rumA*::*CAT)* mutant on 3-AT plate, in spite of being *relA*^+^, is due to the overexpression of RelA protein and not due to the slow decay of (p)ppGpp because of the *spoT1* mutation, as growth of KP31 mutant on 3-AT plate is same as that of MG1655 (Fig.2C). Our observation is supported by similar results reported by (Sanchez-Vazquez *et al*., 2019) which also matches the inference of the results described below on growth phenotype of transformants of RelA-CTD (see below).

**Fig. 4:**
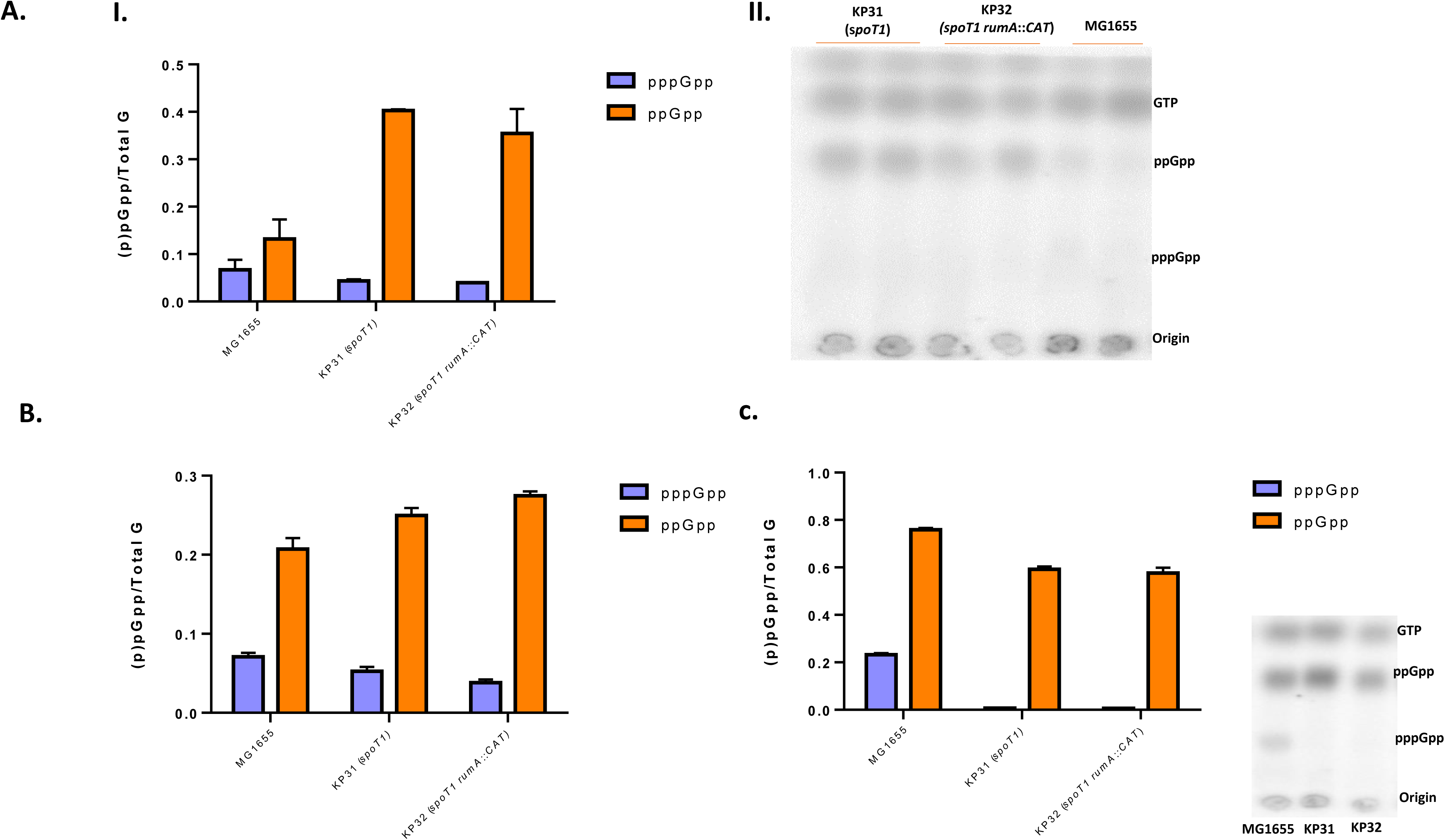
Basal level of (p)ppGpp measurement in KP32 (*spoT1 rumA::CAT*) mutant grown in MOPS glucose at (**A**) 37°C; (**B**) 30°C and (**C**) in response to isoleucine starvation.

### Low to moderate level expression of the plasmid-borne *relA* gene in the *spoT1* mutant recapitulates the inhibitory growth phenotype of *rumA*::*CAT* mutant

We performed the experiment with *relA* gene cloned in two types of plasmid vectors. (i) The *relA* gene in pGB2*-relA* (a kind gift from M. Cashel) is expressed from *lac* promoter in the low copy pSC101 replicon, pGB2 (Churchward *et al*., 1984). The expression of the RelA from this plasmid was found to be partially de-repressed in MG1655 and its *spoT1* mutant (KP31) (Fig.5A(III)), although not to a level as high as in KP32 (*spoT1 rumA*::*CAT*) mutant. Nevertheless, the growth phenotype of the pGB2-*relA* transformants of *spoT1* strain (KP31) was identical to that of KP32; severely inhibited in minimal glucose medium even in the absence of IPTG supplementation (Fig.5A(Ia, Ib)). Similarly, the unusual stringent response of KP31 (*spoT1*)/pGB2 *relA* is indistinguishable from that of the KP32 mutant (Fig. 5A (Ic)), exhibiting strikingly retarded growth on minimal agar plate supplemented with 3-AT even in the absence of IPTG. Importantly, levels of (p)ppGpp did not differ detectably between transformants of KP31 (*spoT1*) /pGB2 and KP31/pGB2-*relA* (Fig.5A (IIa, IIb)), even though the RelA protein is elevated in the latter, and the levels were same as in KP31 (*spoT1*) and KP32 (*spoT1 rumA*::*CAT*) mutant (Fig.5A(III)).

(ii) The *relA* gene in the medium copy plasmid pTE6 (pBAD18 KAN *relA*) is expressed from its own promoters’ present ∼800 bp in the upstream DNA of *rumA* gene (Fig.1) (Brown *et al*., 2014, Metzger *et al*., 1988, Nakagawa *et al*., 2006). Here too, expectedly, the growth of pTE6 transformants of KP31 (*spoT1*) was drastically impaired in relation to those of MG1655, more acutely in minimal medium (Fig.5B (Ia, Ib)) than in LB (data not shown). Furthermore, the stringent response of the *spoT1* transformants is abnormal, similar to that of KP32 (*spoT1 rumA*::*CAT*) mutant, in that the poor growth on amino acid starvation plate (minimal agar plate supplemented with 3-AT) is almost same as that on minimal agar (Fig.5B (Ic)). The result clearly indicates that the phenotype of KP32 (*spoT1 rumA::CAT)* could be also reproduced by moderate level expression of *relA* gene using the second plasmid system, although the possibility of levels of (p)ppGpp being higher in the *spoT1* mutant which may reduce the growth, cannot be ruled out.
(iii) The plasmid pALS10 (a gift from M. Cashel) extensively used for ectopic production of (p)ppGpp expresses the RelA protein under the control of the *tac* promoter in *colE1* replicon (Schreiber *et al*., 1991). The growth phenotype of the transformants of *spoT1* mutant merits attention (Fig.5C). The colonies were small, sick and extremely slow growing on LA agar in the absence of induction of RelA by IPTG, and fail to regrow. The plasmid also affects the growth of MG1655 strain appreciably in comparison to pALS14 (containing the inactive *relA* gene) (Fig.5C), the phenotype described for long in the literature (Schreiber *et al*., 1991, Svitil *et al*., 1993). We believe that the retardation of the growth of MG1655 by multiple copies of *relA* is independent of (p)ppGpp synthesis, at least partially, for following reasons - (i) pALS10 plasmid contains the gene for *lacI* repressor for increasing the host range of the plasmid and for regulated expression of *relA* so that the basal level of expression of the *relA* is not toxic in multiple copies. Indeed, the (p)ppGpp levels in the cells of MG1655 carrying pALS10 or its equivalent (Mechold *et al*., 1996, Schreiber *et al*., 1991, Svitil *et al*., 1993, Zhu & Dai, 2019) grown in the medium lacking IPTG is comparable to the cells containing pALS14 even though the un-induced levels of RelA protein are significantly high (Schreiber *et al*., 1991). This observation is the basis of the authors’ (Sanchez-Vazquez *et al*., 2019) surmising that the cellular pool size of (p)ppGpp is dictated by the *relA/spoT* genotype of the host and is independent of the basal level expression of the cloned copy number of *relA*. (ii) A recent finding by Turnbull *et al*., (2019) demonstrates a strong inhibition of the growth by high levels expression of the C-terminal fragment of RelA supporting the contention that the inhibitory effects of RelA overexpression may involve (p)ppGpp-independent component as well. The steady state pool size of (p)ppGpp in KP31 (*spoT1)*/pALS10 cannot be estimated due the lethality of the plasmid.

**Fig. 5:**
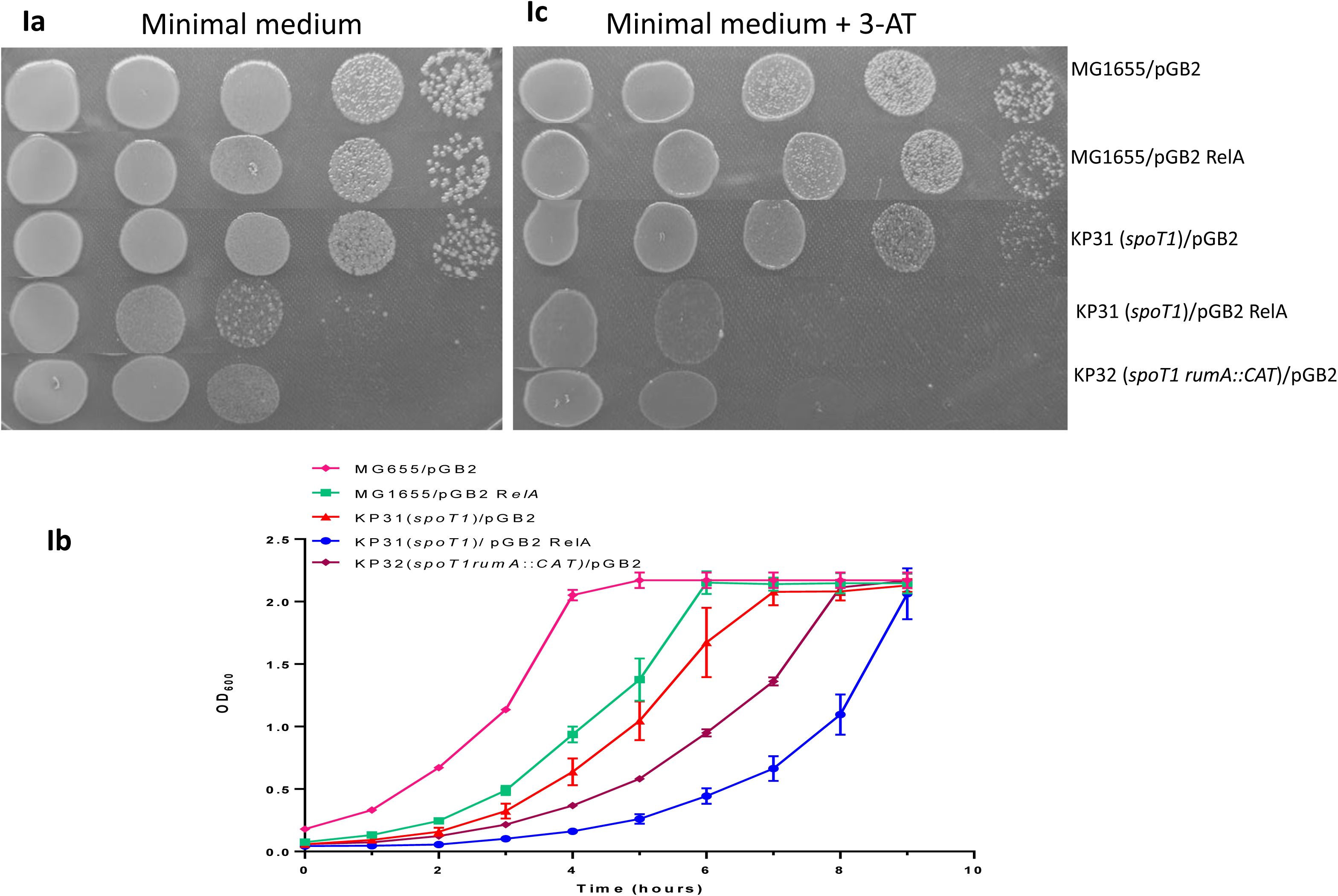
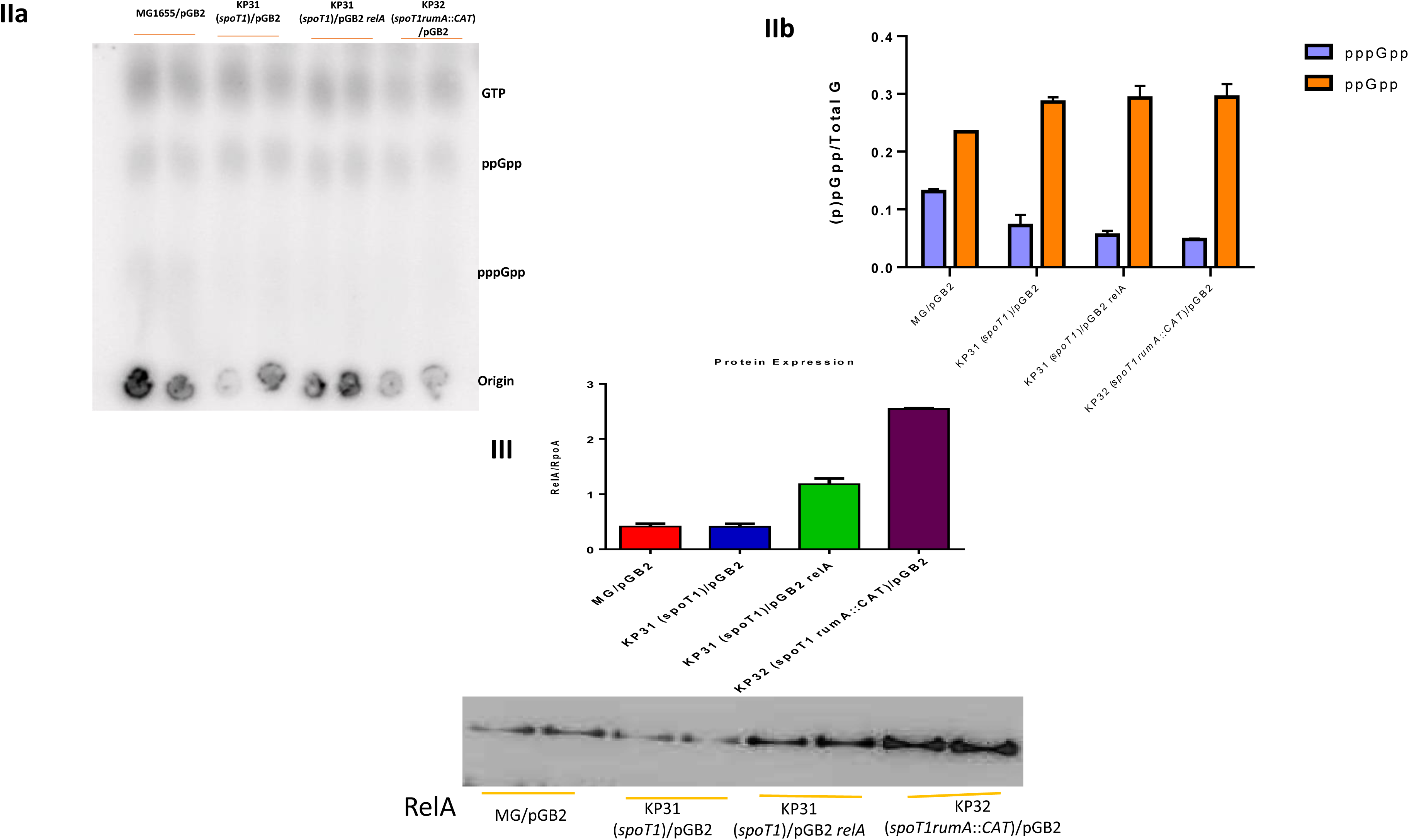
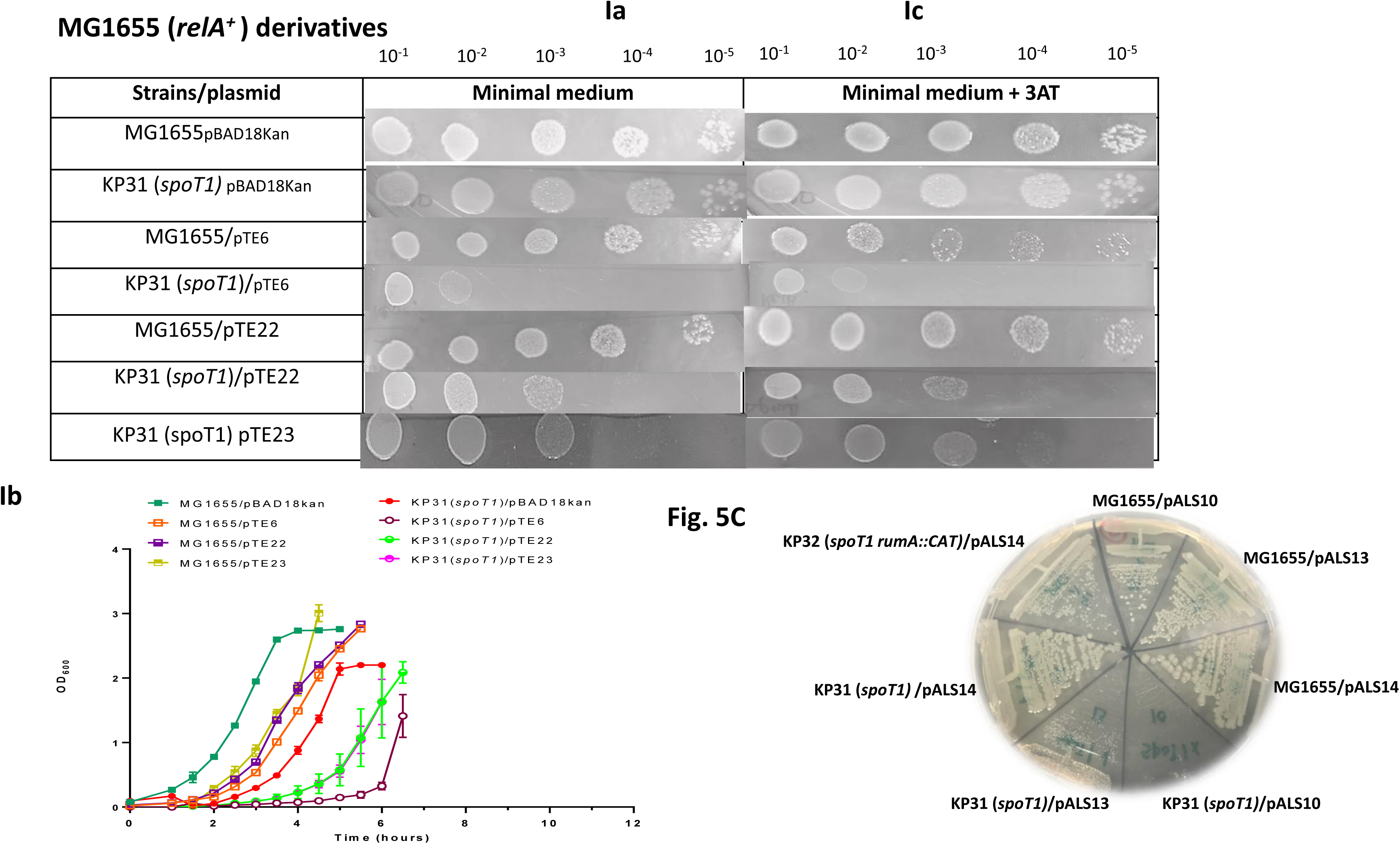
Plasmid mediated low to moderate level expression of RelA in the *spoT1* mutant background (KP31) recapitulates the growth phenotypes of *rumA*::*CAT spoT1* mutant, KP32: 5A: **(Ia)** Growth of pGB2 *relA* transformants of WT and *spoT1* mutant on minimal agar plate at 30°C, **(Ib)** Growth curve of the pGB2 and pGB2 RelA transformants of *spoT1* and *spoT1 rumA::CAT* mutants was measured in minimal broth at 37^0^C. The growth curves were plotted from three independent experiments, **(Ic)** Growth of pGB2 *relA* transformants of WT and *spoT1* mutant on minimal agar plate supplemented with 3-ATat 37°C. **(IIa, IIb)** TLC measurement and quantitation respectively of basal level of (p)ppGpp in KP32 (*spoT1 rumA::CAT*), MG1655/pGB2 *relA* and KP31 (*spoT1*)/ pGB2 *relA* in cells grown in MOPS glucose medium, **(III)** Quantification of RelA protein in KP32, MG1655/ pGB2 *relA* and *spoT1* mutant (KP31)/ pGB2 *relA* by Western blot; (5B): Moderate level expression of RelA-CTD (pTE22 (RelA-CTD1)) and (pTE23 (RelA-CTD2)) selectively inhibits growth of the *spoT1* mutant: (Ia) Growth phenotype of *relA*^+^ strains, MG1655 and KP31 (*spoT1*), transformed with following plasmids - control (pBAD18Kan), pTE6 (pBAD18Kan RelA), pTE22 (RelA-CTD1) and pTE23 (RelA-CTD2) on minimal medium and minimal medium supplemented with 3-AT; **(Ib)** growth curve measurement of the same set of strains as in Ia in minimal broth at 37^0^C. The growth curves were plotted from three independent experiments; **(5C)** Severe growth retardation phenotype of transformants of KP31 (*spoT1*)/pALS10 on LA + Amp plate. The plate was incubated at 30°C.

### Overexpression of fusion of N-terminal (1-119 amino acids) with RelA-CTD (454-744 and 406-744) renders growth inhibition by RelA-CTD, (p)ppGpp dependent

Our observation that the increased level of expression of RelA in *rumA::CAT spoT1* mutant (Fig.3 C) or the leaky expression of plasmid copies of *relA* in the absence of the inducer IPTG (Fig.5III) is not associated with a corresponding increase in the (p)ppGpp levels (Fig4; Fig.5A (IIa, IIb)), is not without precedence (Sanchez-Vazquez *et al*., 2019). This observation applied to both wild type (Schreiber *et al*., 1991, Gropp *et al*., 2001) and to the *spoT1* mutant (Fig 4; Fig.5A (IIa, IIb)), albeit the level of (p)ppGpp is characteristically higher in the *spoT1* mutant than in the wild type and independent of *rumA::CAT* mutation in KP32 or presence of additional copies of plasmid encoded *relA* in KP31/pGB2 *relA* (Fig.4II, Fig.5 (IIb)). Strikingly, the growth retardation of the *spoT1* mutant containing multiple copies of RelA is more severe than that of the *spoT1* mutant itself, even though the levels of (p)ppGpp are the same in each of the two conditions (Fig. 2A, 2B, 2C, 5A (Ia, Ib), 5B (Ia, Ib), 5C).

The result led us to assume that the RelA’s (p)ppGpp synthesis function may not be involved in the *spoT1* mutation-dependent growth retardation phenotype. It is thus reasonable to test the growth effects of RelA-CTD in the *spoT1* mutant. As we understand, the N-terminal domain (1-390) of the RelA protein consists of 1-200 amino acids constituting inactive hydrolytic (HD) domain and 201-390 amino acids comprising (p)ppGpp synthesis domain (SYN). Thus, presuming the N-terminal 119 amino acids to be devoid of any discernible function, the simple strategy of cloning CTD involved removing sequences internal to the two *Pvu*II sites at positions 354 and 1362 in the *relA* gene. This fortuitous cloning uncovered an unexpected layer of regulation of *relA*.

Two types of in-frame fusion of N-Terminal 119 amino acids with RelA-CTD were constructed starting with the plasmid pTE6 (pBAD18 Kan *relA*). (i) The plasmid construct pTE22 represents the N-terminal 119 amino acids fused to the C-terminal domain comprising of 455-744 amino acids, generated by *Pvu*II digestion and intramolecular ligation of the *relA* DNA in the plasmid pTE6 (pBAD18 KAN*relA*). The truncated RelA is insufficient in complementing the stringent response defect of *relA1* mutation in MC4100 (Fig.S2 B). The fact that substitution of G251E by site-directed mutagenesis results in inactive mutant protein that has lost (p)ppGpp synthesis ability (Gropp *et al*., 2001), and that the *Pvu*II deletion removes the critical G251 amino acid important for (p)ppGpp synthesis, reaffirms the functional inability of the truncated protein to synthesise (p)ppGpp.

(ii) The plasmid construct pTE23 represents in-frame fusion of N-terminal 119 amino acids with the 406-744 portion of RelA-CTD (Turnbull *et al*., 2019), generated by PCR (described in Materials and Methods). An additional, intact TGS domain (200-396) in pTE23 however, is not required for binding to the ribosomes (Gropp *et al*., 2001, Turnbull *et al*., 2019).

Remarkably, the growth of the transformants of *spoT1* mutant with each of the two plasmids (pTE22, pTE23) was indistinguishable; retarded to the same extent on both minimal agar containing glucose as the sole carbon source and on minimal medium supplemented with 3-AT but less than that by the full length *relA* (pTE6) whereas transformants of MG1655 were unaffected for growth (Fig.5B (Ia, Ib, Ic)). The severity of the effect on growth in minimal medium is evidently due to (p)ppGpp levels being high in minimal glucose medium (nutrient poor) than in LB.

### Abrogation of the reduced growth effect of moderate expression of RelA-CTD by *relA1* mutation is reversed by *spoT202* mutation

The result that growth of the *spoT1* mutant is retarded by moderate level expression of RelA-CTD in two fusion constructs, pTE22 (1-119 NTD fused to 406-744 CTD) and pTE23 (1-119 NTD fused to 454-744 CTD), which is without any effect in the wild type strain (Fig. 5B (Ia, Ib)) suggests that the higher basal levels of (p)ppGpp in the *spoT1* mutant may be required for manifestation of the growth effects. In agreement with this assumption, the *relA1* mutation, which lowers (p)ppGpp levels in the *spoT1* mutant, also abolishes the *spoT1*-specific effects of growth by the RelA-CTD (Fig.6 (Ia, Ib)). If the growth effects of RelA-CTD are indeed accentuated by higher basal levels of intracellular levels of (p)ppGpp, it should be possible to restore slow growth phenotype of the RelA-CTD in *spoT* mutants that contain levels of (p)ppGpp higher than in the *relA1 spoT1* mutant. Mutant *spoT* alleles, *spoT202, −203, −204* that are increasingly compromised for their (p)ppGpp hydrolysis function have been described to reverse the 3-AT sensitive phenotype of *relA1* mutants as a result of the enhanced (p)ppGpp levels (Sarubbi *et al*., 1988),Fig.S4 B), and accordingly exhibit slow growth (Fig.S4 A). Because of the increasing severity of the *spoT* alleles on growth, we chose to work with *spoT202* mutation, which is more severe than *spoT1* in its ability to elevate (p)ppGpp levels but less than that by *spoT203/204* (Sarubbi *et al*., 1988), KP56 (*relA1 spoT202 zib563*::Tn*10*) and its isogenic derivative, KP57 (*relA1 zib563*::Tn*10*) were constructed by P1-mediated cotransduction of *spoT202* mutation with *zib563*::Tn*10* (Table 1). The control vector, pBAD18Kan, pTE22 (RelA-CTD1) and pTE23 (RelA-CTD2) were each transformed in the isogenic pair and the growth was measured qualitatively (Fig.6II A) and quantitatively (Fig.6II B). The phenotype of effect of overexpression of fusion RelA-CTD on growth was largely reproduced in the *spoT202* mutant, in that the pTE22 (RelA-CTD1) and pTE23 (RelA-CTD2) transformants of KP56 (*relA1 spoT202*) were appreciably slow growing in comparison to the control vector pBAD18Kan transformants (Fig 6 (IIa, IIb)), strongly indicative of the regulation of C-terminal function by (p)ppGpp. The longer lag in the growth of KP31/ (pTE22/pTE23) transformants (Fig. 5B (Ib)) in comparison to that of KP56 /pTE22 (Fig.6II B) might be due to the potential for dynamic changes in the concentration of (p)ppGpp effected by RelA in the nutrient poor minimal medium in the former strain than in the latter. Furthermore, the growth of KP58 (*spoT202 relA^+^*) transformants carrying pTE22 or pTE23 was severely reduced in comparison to that of pBAD18kan plasmid transformants (Fig.6C (I, II)). It must be noted here that, unlike the observation of (Sarubbi *et al*., 1988) which describes an inability of *spoT 202/203* mutation to coexist with *relA*^+^, we were able to transduce each of the *spoT* mutation into the wild type MG1655 strain. As expected, growth of *relA^+^ spoT 202* transductants, KP58, was drastically slow when compared to that of *relA1 spoT202* mutants (Fig.S4A). Growth of KP56 (*relA1 spoT202*)/pTE22 (RelA-CTD1) transformants on minimal agar plate supplemented with 3-AT was similarly compromised/reduced to the same extent as on minimal agar (data not shown). These results convincingly demonstrate that the growth effects of multicopy expression of RelA-CTD are indeed potentiated by higher levels of intracellular (p)ppGpp and that the enhanced levels of (p)ppGpp effected either by *spoT1* or *relA1spoT202* mutations are sufficient for the manifestation of the growth retardation/abnormal stringent response phenotypes. Given the known function of the C-terminal domain of RelA to bind ribosomes (Gropp et al., 2001), our results seem to suggest that this binding is modulated by the levels of (p)ppGpp in the cell; and since RelA-CTD binding to ribosomes has been shown to slow translation rates (Turnbull *et al*., 2019) the effect on the growth can be rationalized.

**Fig. 6:**
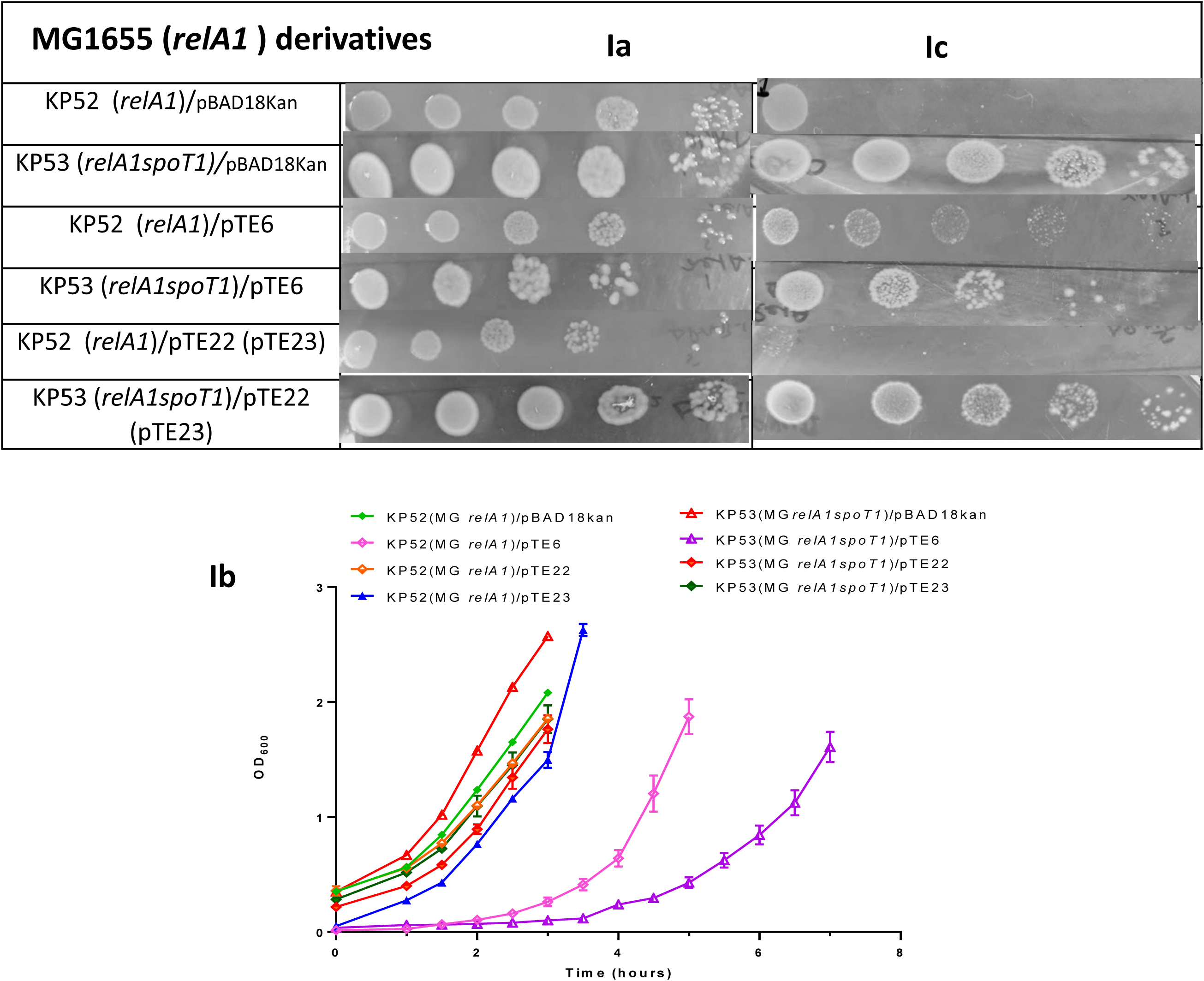
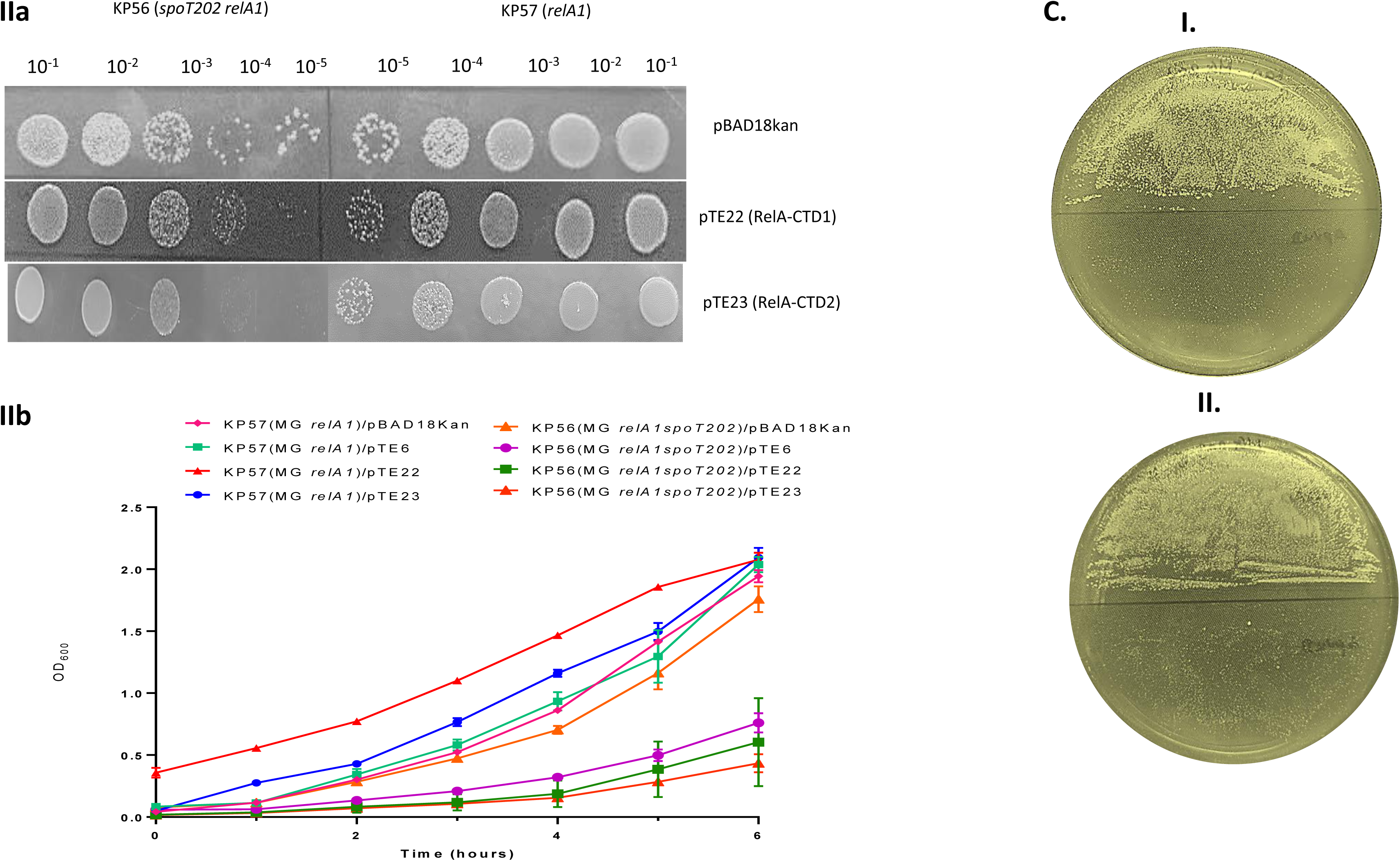
Obliteration of the *relA1* mutation-mediated growth inhibitory effects of overexpression of RelA-CTD in *spoT1* mutant is reversed by *spoT202* mutation. **(Ia)** Growth on minimal medium and minimal medium supplemented with 3-AT of *relA1* mutant strains KP52 (MG1655 *relA1*) and KP53 (MG1655 *relA1spoT1*) transformed with plasmids - control (pBAD18Kan), pTE6 (pBAD18Kan RelA), pTE22 (RelA-CTD1) and pTE23 (RelA-CTD2); **(Ib)** Growth curve of transformants of same set of strains as in Ia was measured in Minimal broth at 37°C. The growth curves were plotted from three independent experiments. **(IIa)** Growth phenotype of pTE22 (RelA-CTD1) and pTE23 (RelA-CTD2) transformants of KP56 (*relA1 spoT202*) is significantly retarded in comparison to that of control plasmid, pBAD18Kan and the CTD plasmid transformants of KP57. **(IIb)** Growth curve measurements of the pTE22, pTE23 plasmid transformants of KP56 and KP57. The growth curves were plotted from three independent experiments. (**C)**: Severe growth effects of overexpression of RelA-CTD1 (pTE22) **(I)**, and pTE23 (RelA-CTD2) **(II)** in *relA^+^ spoT202* mutant KT58. The growth of the pBAD18Kan transformants of each of the mutant was obvious on the plates within 16 hrs. of incubation in contrast to that of pTE22/pTE23 transformants. The plates were photographed after 24 hrs of incubation at 37°C.

**Table 1:**
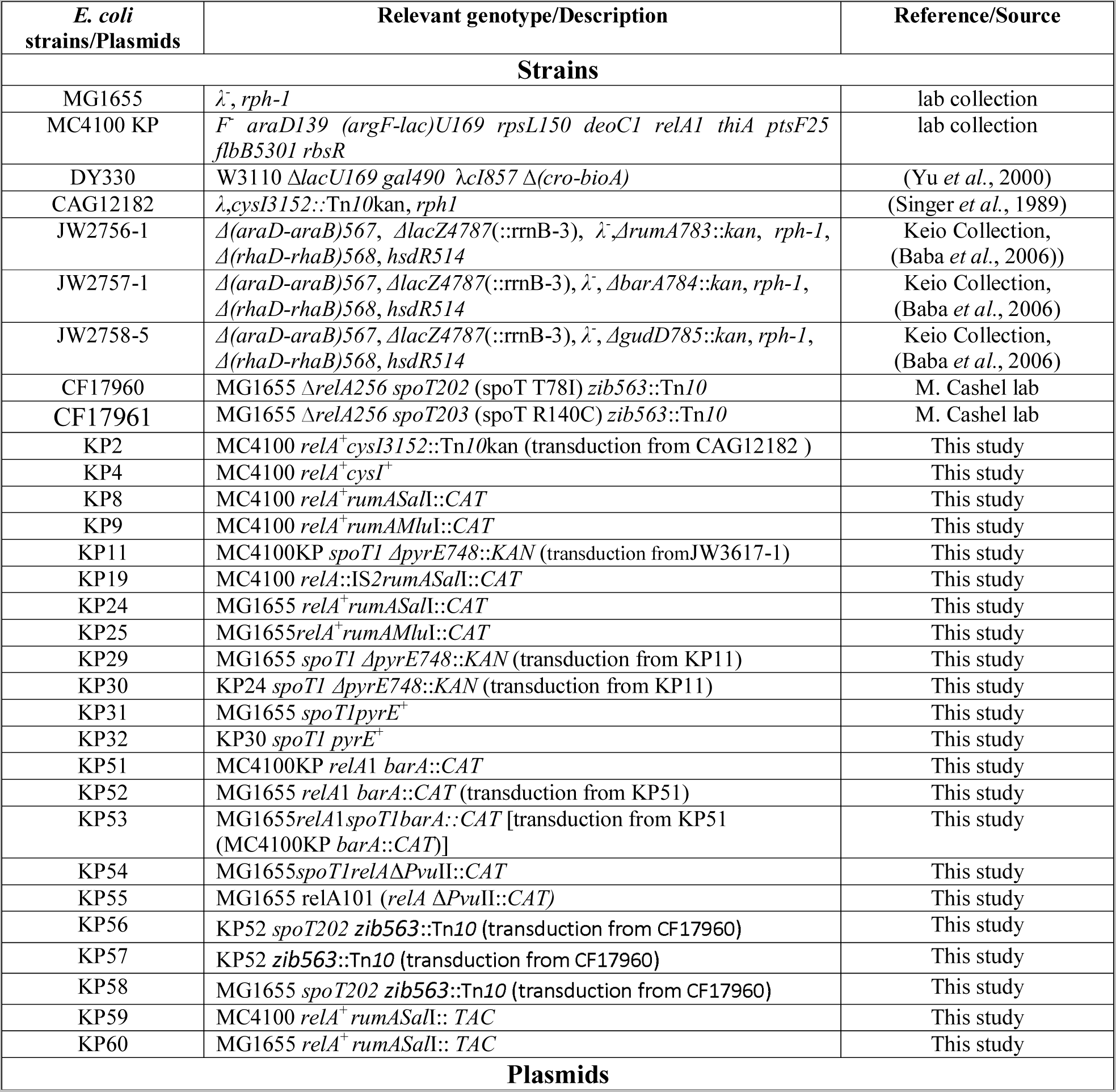
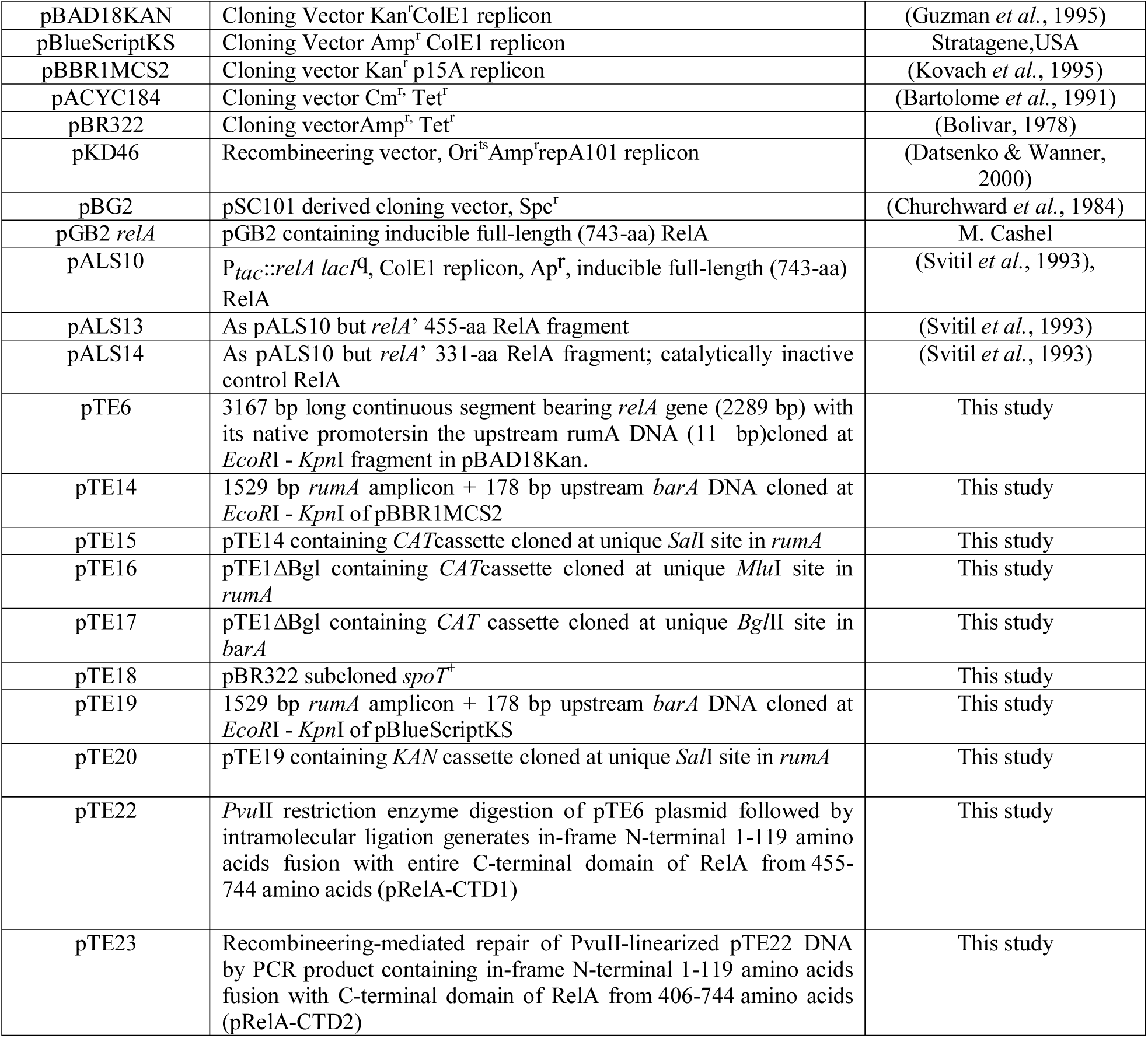
Strains and plasmids used in this study.

In summary, the following results likely contribute to further our understanding of the regulation of RelA and its (p)ppGpp synthesis function. Sustained low to moderate level of expression of RelA in the *spoT1* mutant was associated with phenotypic consequence of slow growth, clearly absent in the wild type strain, suggesting that higher than the normal basal level of (p)ppGpp is required for the manifestation of the growth phenotype. This is further validated by reproduction of a large part of the growth retardation effect by expression of RelA-CTD (Fig. 5B (Ia, Ib)), and abolition of the slow growth phenotype by decreasing the intracellular levels of (p)ppGpp in the *spoT1* mutant by *relA1* mutation (Fig.6A (Ia, Ib)). The requirement of higher levels of (p)ppGpp for RelA-CTD mediated inhibition of growth can be also met by substitution of chromosomal RelA’s (p)ppGpp synthesis function with *spoT202* mutation which elevates intracellular (p)ppGpp levels in *relA1* mutant due to the an inactive hydrolytic activity of the SpoT protein (Fig.6 (IIa, IIb))(Sarubbi *et al*., 1988). We were able to assign this (p)ppGpp regulatory function to the 119 amino acids NTD domain as its in-frame fusion with RelA-CTD, both 406-744 (RelA-CTD2) and 455-744 (RelA-CTD1) rendered the growth inhibitory effects of its expression dependent on the elevated cellular levels of (p)ppGpp. Our findings may suggest a mechanism that provides a rationale for *in vivo* regulation of RelA’s (p)ppGpp synthesis activity.

## Discussion

Our results seem to unfold a new aspect of the regulation of RelA demonstrating that the aberrant stoichiometry of RelA (more precisely RelA-CTD) to ribosomes is inhibitory to growth, the severity of which is determined by an incremental rise in the basal levels of (p)ppGpp, therefore independent of its synthesis by RelA. This inference is apparent from our unanticipated result that mutational and/or growth condition mediated elevation of (p)ppGpp pool size potentiates inhibition of growth by moderate level expression of RelA-CTD. Expression of in-frame fusion of the N-terminal 119 amino acids of the HD domain with the RelA-CTD (406-744, containing TGS domain) or (454-744, lacking the TGS domain) was equally effective in causing growth inhibition of *spoT1* mutant, but not that of wild type. Our results are compatible with the model that the growth inhibitory effect of RelA-CTD is due to its interaction with the target ribosomes which is modulated by (p)ppGpp levels in the cell, the toxicity effects, are nevertheless independent of the (p)ppGpp synthesis function of RelA.

It must be noted that, unlike the growth inhibitory effect of high-level expression of RelA-CTD, the growth phenotypes we describe here are more pertinent and relevant to physiological concentration of RelA because the protein is expressed from its own promoters in a low copy to medium copy vector and translated from its own ribosome binding site (SD sequence) not optimised for very high-level expression (Turnbull *et al*., 2019). Importantly, the high-level multicopy expression of RelA-CTD^454-744^ (a gift from M Roghanian) inhibits growth of strains irrespective of their genotype; on the contrary, expression of the same RelA-CTD gene in a low copy plasmid, pNDM220:*relA*^CTD^ (a gift from M Roghanian) does not affect either the growth of wild type or its *spoT1* derivative (data not shown) or interfere with the expression of stringent response (Turnbull *et al*., 2019), proving that NTD’s HD domain contains features for (p)ppGpp-dependent regulation of growth effects of RelA-CTD.

Some of the observations pertinent to our studies regarding the ectopic expression of plasmid-borne relA from a controlled promoter is the lack of proportional increase in the (p)ppGpp levels and the extent of growth inhibition induced by overexpression of full length RelA (Schreiber *et al*., 1991, Gropp *et al*., 2001), explained earlier as being due to toxicity of full length RelA protein (Schreiber *et al*., 1991). The implication of this and the results presented here is that the growth inhibitory effects of RelA are due to (i) the product, (p)ppGpp-mediated inhibition of transcription of stable RNA and ribosomal protein synthesis and (ii) (p)ppGpp-responsive altered binding of RelA-CTD to ribosomes, slowing rate of protein synthesis. In the recent study, Sanchez-Vazquez *et al*., (2019) concluded that the leaky basal level expression of active RelA does not alter the (p)ppGpp levels in the cells which are rather dictated by the chromosomal status of *relA/ spoT* genes, which strikingly corroborate ours (Fig. 4, Fig 5A (IIa, IIb)). Our finding that growth inhibition effect of leaky, un-induced, low (Fig. 5A (Ia, Ib)) to moderate level expression of RelA is more acute in the *spoT1* mutant than in the wild type strain and that the un-induced multiple copy expression is lethal in the *spoT1* mutant (Fig.5C), merits investigation.

How relevant is our observation? The positive activation of RelA by its own product, (p)ppGpp (Shyp *et al*., 2012), explains the rapid increase in the concentration of (p)ppGpp in the cell in response to the signal of amino acid starvation, mandating a negative component of regulation of its synthesis as well. When the concentration of (p)ppGpp reach the threshold required for a global effect on transcriptome and proteome, its cascading effect must be countered/tempered by a negative regulation of RelA’s synthesis activity. One of the proposed negative means of regulation of RelA activity is a decrease in the (p)ppGpp levels affected by restoration of the amino acid pool under favourable growth condition. It makes sense that the mechanism proposed here - the RelA-CTD interaction with the target ribosome is modulated by incremental rise in (p)ppGpp which possibly inhibits its synthesis by the N-terminal, complements the negative regulation. This continuous responsiveness of RelA function to (p)ppGpp levels in the cell could likely be an *in vivo* mechanism of negative modulation of (p)ppGpp synthesis. The *in vitro* observation that (p)ppGpp synthesis is maximum at less than 10:1 of 70S:RelA ratio and the synthesis decreases as the stoichiometry increases (Wendrich *et al*., 2002, Kudrin *et al*., 2018) seems to support our finding that moderate level of RelA overexpression does not lead to (p)ppGpp synthesis *in vivo* (Schreiber *et al*., 1991, Gropp *et al*., 2001).

The possibility is worth considering that the increase in the levels of RelA in the exponentially growing cells slow the growth by affecting the higher affinity of EFTu.tRNA.GTP ternary complex at the A site on the ribosome (Winther *et al*., 2018, Loveland *et al*., 2016).

Interestingly, results of Syal *et al*., (2015) point to similar conclusions. They showed that Rel-CTD of *M*. *smegmatis* binds to (p)ppGpp and suggest this binding to be important for providing a negative feedback mechanism of regulation to adjust the (p)ppGpp levels in the cell. They confirmed that mutations in the Rel protein implicated in the binding of (p)ppGpp accordingly increased p(p)ppGpp synthesis and reduced hydrolytic activity.

This work raises the question - has RelA’s catalytically inactive HD got any role or function for the protein? The evidence is largely tantalizing. On the lines that there is reciprocal inter-domain regulation between HD and the SYN domains of the NTD of long RSHs (Tamman *et al*., 2019, Takada *et al*., 2021), it is interesting to note that mutations in the inactive HD domain of E. coli RelA reducing the SYN activity (Montero *et al*., 2014, Spira & Ospino, 2020) may reflect the relic of this regulation. Although the catalytic amino acid residues of the HD domain are clearly absent in the monofunctional RelA, there is evolutionary conservation of the HD sequence all along this region between bifunctional RSHs and RelA, suggesting a possible important function. Recently, Takada *et al*., (2021) have clearly demonstrated the localization of (p)ppGpp binding site in the NTD domain of RelA protein not involving the catalytic SYN site. In light of this, our result that the (p)ppGpp mediated regulatory function is present in the NTD’s HD domain for regulating the growth inhibitory function of RelA-CTD may not seem implausible.

## Materials and Methods

### Growths Conditions

Bacterial cells were normally grown in Luria Bertani (LB) broth or in M9 minimal broth (MB) at 37°C or 30°C with shaking. 2×YT broth was used for bacterial cell growth during electroporation. When necessary, media were supplemented with Kanamycin (50µg/ml), Ampicillin (100µg/ml), Chloramphenicol (12µg/ml), Tetracycline (10µg/ml), Streptomycin (100µg/ml), Spectinomycin (50µg/ml). Isopropyl β-D-thiogalactopyranoside(IPTG) was used at a final concentration of 1mM. DNA manipulations were carried out according to protocols described in (Sambrook, 1989). P1 transduction was carried out by method described by (Miller, 1992). Strains used in this study are the derivatives of *E. coli* K-12. Strains and plasmids are listed in Table 1. Primers used in this study are listed in Table 2.

**Table 2:**
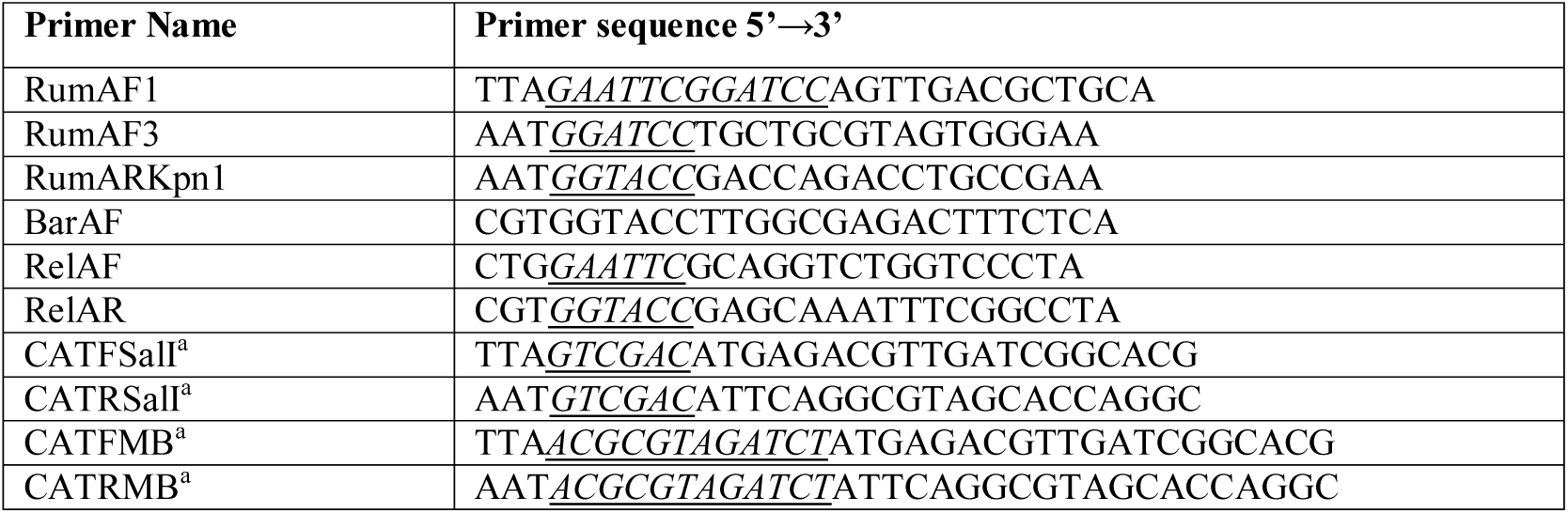
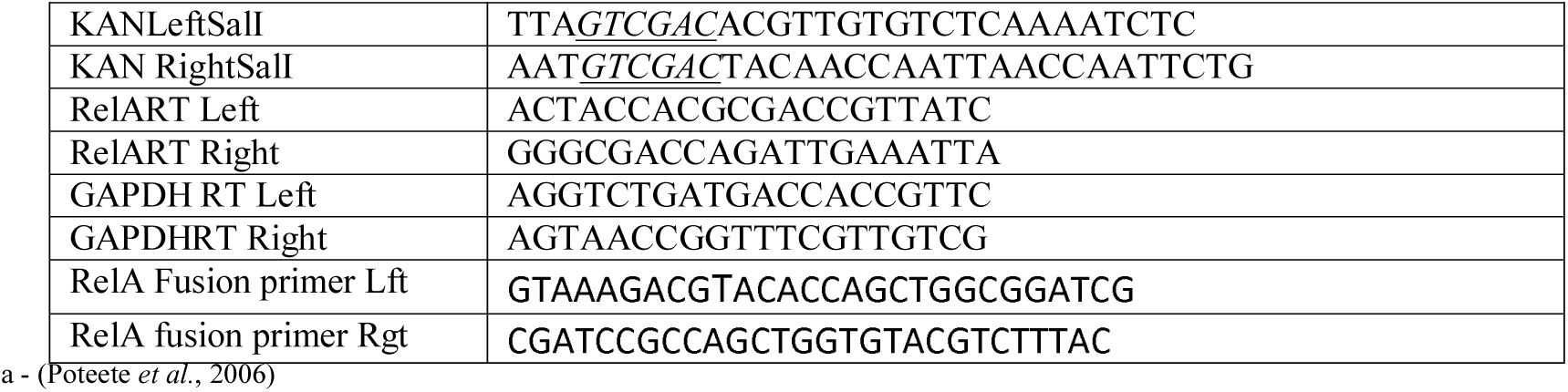
List of Primers used in this work.

## Methods

### Insertion of *CAT/KAN* cassette in chromosomal *rumA* gene

The *rumA* amplicon (1529 bp), generated using RumAF1-RumARKpnI primers, was cloned in to the broad host range vector pBBR1MCS2 (Kovach *et al*., 1995) and vector pBluescriptKS at *EcoR*I-*Kpn*I sites yielding pTE14 and pTE19 respectively.

1. Amplification of 824 bp *CAT* gene from pACYC184 vector (Bartolome *et al*., 1991) was carried out by 2 pairs of CAT primers. (i) CATFSalI and CATRSalI primers were used for cloning *CAT* amplicon into pTE14 vector at unique *Sal*I site to generate pTE15, (ii) Using CATFMB and CATRMB primer pair, *CAT* amplicon was inserted at *Mlu*I site in *rumA* to generate pTE16 construct.
2. Amplification of 944 bp *KAN* gene from pBAD18Kan Vector (Guzman *et al*., 1995) was carried out by KAN primers KANLeftSalI and KANRightSalI and cloned at unique *Sal*I site of pTE19 to generate pTE20. Insertion of *CAT*/*KAN* cassette at the plasmid level was confirmed by restriction digestion.

We replaced chromosomal *rumA* gene with the mutant allele by recombineering in *E. coli* strain DY330 (Yu *et al*., 2000) and also using the portable pKD46 plasmid (Datsenko & Wanner, 2000) in MG1655 strain. The linear PCR DNA of mutant *rumA* gene containing *CAT* (2335 bp) or *KAN* cassette (2455 bp) was amplified using RumAF1-RumARKpnI primers and the plasmids pTE15 and pTE20 used as templates. The PCR DNAs were purified by gel elution method and transformed into electrocompetent cells. The genomic mutations in *rumA* containing insertion of *CAT*/*KAN* in each case were transduced into a fresh background of MC4100 and MG1655 through P1*kc*-mediated generalized transduction as described by (Miller, 1992). The orientation of antibiotic cassette was determined by PCR using antibiotic cassette specific forward primer in combination with *rumA* gene specific forward and reverse primers (data not shown). In the transduced MC4100 derivatives, the *relA*^+^ gene was also confirmed by PCR and sequencing (data not shown). Similarly, the P1 transduction-mediated transfer of *spoT1* mutation in MG1655 was confirmed by DNA sequencing of the representative strains.

### Construction of KP4 (MC4100 *relA*^+^)

The strains used in the study were constructed by P1 transduction. *relA*^+^ allele was cotransduced into MC4100KP (*relA1 spoT1*) with *cysI3152::*Tn*10*kan (CAG12182). For markerless strain construction in this cross, CysI^-^ transductants were converted into CysI^+^ prototrophs, again by P1 transduction, screening for loss of Kan^r^ gene. The *relA* gene structure was confirmed by PCR using RelAF - RelAR primers. *relA1* gene of MC4100 generates 3.561 Kb amplicon due to an IS2 (1.327 Kb) insertion in the *relA* whereas the size of the wild type *relA* gene is 2.234 Kb (data not shown).

### Exchange of *spoT* alleles from MC4100 to MG1655

The *spoT1* allele of MC4100KP was first linked to *ΔpyrE748*::*KAN* (KP11) marker (∼67% cotransduction), followed by its transfer into MG1655 and KP24 through P1 transduction generating KP29 (*spoT1 ΔpyrE748*::*KAN*) and KP30 (*rumASal*I::*CAT spoT1 ΔpyrE748*::*KAN*) respectively; this is followed by selection of *pyrE^+^* derivatives, KP31 (*spoT1*) and KP32 (*spoT1 rumASal*I::*CAT*) on minimal medium in the absence of uracil.

### Assessment of *relA* phenotypes

The growth phenotypes of RelA^+^ and RelA^-^ were scored on one of the three types of plate as described below. (i) M9 minimal medium supplemented with glucose (0.2%) and all amino acids except histidine (4µg/ml), adenine (1 mM), thiamine (1 mM), and 3-AT (15 mM)(Silva & Benitez, 2006). (ii) M9 glucose medium containing serine, methionine, glycine, (SMG) (100 µg/ml each), adenine (50 µg/ml), thymine (50 µg/ml), and calcium pantothenate (1 µg/ml) (Silva & Benitez, 2006). (iii) Amino acid starvation was induced by addition of 0.5-1mg/ml serine hydroxamate in minimal medium containing 0.2% glucose (Tosa & Pizer, 1971).

### Determining growth

Growth curve experiments were performed in either LB broth and/or M9 Minimal broth. Overnight grown cultures were diluted 1:100 in 50ml fresh media, shaken at 37°C temperature, unless specified otherwise. OD_600_ was monitored. Growth rate experiments were repeated thrice.

### Isolation of *spoT*^+^ gene in pBR322

A MG1655 genomic DNA library, prepared by partial digestion of chromosomal DNA with *Sau3A*I, was cloned into the *BamH*I site of pBR322. P1 lysate made on the library of pool of 10,000 independent clones was transduced into KP32 (*rumASal*I::*CAT*) with selection for plasmid-encoded ampicillin resistance on minimal ampicillin agar plate. Transductants were screened for faster growing colonies. Plasmids were prepared from 3 independent colonies that permitted fast growth of KP32 on minimal medium and restored growth on minimal medium + 19 amino acids lacking His supplemented with 3-AT (normal *relA*^+^ phenotype).

### Quantitation of RelA Protein expression by Western blot

Cultures in mid-exponential phase at an OD_600_ of 0.6. The cells were centrifuged and the pellet was resuspended in Laemmli buffer and sonicated. An estimated 20 µg protein was loaded on a 10% SDS-Polyacrylamide gel and then electrotransferred to Nitrocellulose membrane (Bio-Rad). The blots were probed overnight with the monoclonal anti-RelA primary antibody (1:2000; Santa Cruz) and anti-β-Galactosidase primary antibody (1:5000; Novusbio) at 4°C. Anti-mouse/rabbit IgG conjugated with HRP (1:2500) was used as secondary antibody. Finally, membranes were developed and visualized with Enhanced Chemiluminescence Western blotting detection system (Bio-Rad). Images were scanned for densitometric analysis using the Image J software.

### RNA isolation and mRNA expression of *relA* gene by semi-quantitative RT PCR

RNA was isolated from strain of MG1655 derivatives using the TRIzol reagent (Invitrogen). Genomic DNA in the sample was removed with DNAseI kit (Invitrogen). A reverse-transcription into first strand c-DNA reaction was performed using 2 μg RNA with MuLV reverse transcriptase kit in a 20 μL reaction volume (Invitrogen) as per manufacturer’s instructions. cDNA was amplified using gene specific primers (Table 2). GAPDH was used as an internal control and the PCR amplicons were analyzed by electrophoresis on 2.0% agarose gel. Images were scanned for densitometric analysis using the Image J software.

### Measurement of (p)ppGpp levels by thin layer chromatography

Estimation of (p)ppGpp was carried out as described in (Cashel, 1994) and (Fernández-Coll & Cashel, 2019). Briefly, Growth conditions were achieved by continuously shaking 24 well plates in a Thermomixer (Eppendorf) and monitoring OD_600_ in unlabeled parallel cultures in the synergy HT plate reader (Biotek). Overnight cultures were grown in MOPS (3-(N-morpholino) propanesulfonic acid) media with 3 mM sodium phosphate and 0.2% Glucose. Cultures were then diluted into the low phosphate medium (0.2 mM) to give specific activities sufficient for nucleotide labelling with 100 µci of P^32^ in 600 μl of total volume per well and grown till ∼0.4 OD_600_. When necessary valine was added to induce isoleucine starvation. 20µl of samples were collected in a tube containing equal volume of 6M formic acid, then frozen at −20°C. The samples were subjected to three cycles of freeze-thaw and then centrifuged. 5µl of the supernatant was applied on surface of TLC PEI-cellulose (Millipore) and resolved in 1.5M KH2PO4, pH 3.4. The chromatogram was dried at room temperature and the pH front portion containing 32 Pi removed. The autoradiograph was exposed overnight on phosphor screens. Finally, the nucleotide spots were visualized by phosphoimager (Typhoon 9400 imager) and quantified with imageJ. The amounts of ppGpp and pppGpp were normalized to the sum of the pppGpp+ppGpp+GTP (referred as total G) observed in the same sample.

### Construction of in-frame fusion of RelA-CTD with 1-119 amino acids RelA-NTD

The in-frame fusion of 1-119 amino acids of RelA-NTD to 454-744 amino acids of RelA-CTD was carried out by removal of *relA* gene sequence between two *Pvu*II sites at positions 354 and 1362 in the pTE6 (pBAD18kan *relA^+^*) plasmid. In-frame fusion of 1-119 amino acids of RelA-NTD to 406-744 of RelA-CTD was carried out by overlapping PCR in two steps. In the first step, two individual PCRs, one each for the NTD and the CTD, were generated. 1-357 bp PCR product for NTD containing 1-119 amino acids was obtained using the primer pair RelAL and RelAfusionprimerRgt. RelA-CTD PCR product from 1014-2235 bp (406 amino acids) was obtained using primers RelAFusionprimerLFT and RelAR. The PCR products of the NTD and CTD reactions were mixed in 1:1 proportion and used as templates for PCR amplification of the fusion product using primers RelAL and RelAR. The fusion PCR DNA (1578 bp) was used to repair the pTE22 (*relAΔPvu*II) linearized at *Pvu*II site by recombineering in host MG1655/pKD46. The insert DNA from each of the two clones (pTE23) was validated by restriction enzyme digestion and also confirmed by DNA sequencing.

## Acknowledgement

Krishma Tailor received financial support from UGC, Govt. of India. Authors are thankful to M. Cashel and M. Roghanian for their help with plasmids and strains. The authors declare no conflict of interest.

---

**Fig. S1: Measurement of growth of *relA^+^* and *spoT1* mutants of MC4100 and MG1655 strains respectively in minimal broth at 37°C.** Growth curve is representative of three independent experiments.

**Fig. S2: pTE18 (pBR322 *spoT*^+^) plasmid suppresses slow growth and stringent response defect of KP8 [MC4100 (*relA^+^ spoT1 rumA*::*CAT*)]. (A)** Growth on minimal agar and **(B)** on amino acid starvation plate (Minimal agar + 3-AT). The plates were incubated at 37°C.

**Fig. S3: Differential effects of *spoT1* mutation on stringent response in strains MG1655 and MC4100 (A)** *spoT1* allele suppresses the stringent response defect of the *relA1* mutation in KP53 (MG1655 *spoT1relA1)* and promotes its growth on minimal medium supplemented with 3-AT, whereas the growth of the strain in MC4100 containing the same combination of alleles is sensitive to 3-AT; **(B)** complementation of the defective stringent response of MC4100 (*relA1 spoT1*) with the plasmid pTE6 (pBAD18Kan RelA) but not by pTE22 (RelA-CTD1) on minimal plate supplemented with 3-AT.

Fig. S4: Growth Phenotypes of *spoT202* mutant derivatives of *relA*^+^ and *relA1* strains:

**A:** Slow Growth of *spoT202 relA*^+^ strain (KP58) on LA agar plate in comparison to those of *relA1* mutant (KP56). The plates were incubated at 37^0^C and photographed after 16 hrs. of incubation. **B**: The stringent response of *relA1 spoT202* mutant was assessed on minimal agar plate supplemented with 15 mM 3-AT. 1 through 5 are *spoT202 relA1* mutants (KP56 being the representative) that grow on the histidine starvation plate due to high intracellular ppGpp levels and 6 through 8 are three independent isogenic *relA1 spoT*^+^ derivatives (KP57 being the representative) that do not grow.

## Notes

### Competing Interest Statement

The authors have declared no competing interest.

