## Supplementary Figure file for "Fusion of the N-terminal 119 amino acids with the RelA-CTD renders its growth inhibitory effects ppGpp-dependent"

### Slide 1
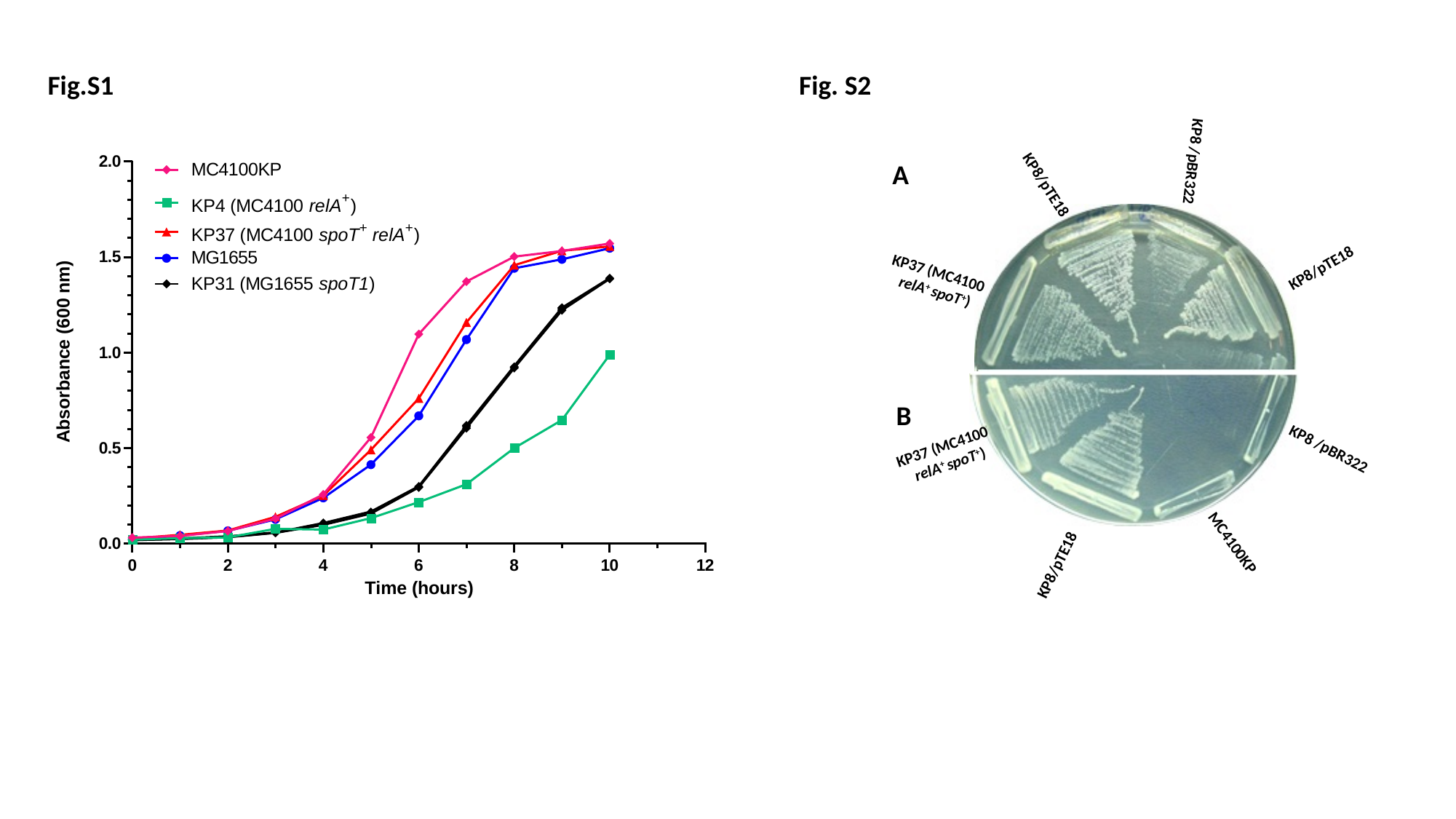

Fig.S1
Fig. S2
KP8 /pBR322
A
KP8/pTE18
KP8/pTE18
KP37 (MC4100
 relA^+^ spoT^+^)
B
KP37 (MC4100
 relA^+^ spoT^+^)
KP8 /pBR322
MC4100KP
KP8/pTE18

### Slide 2
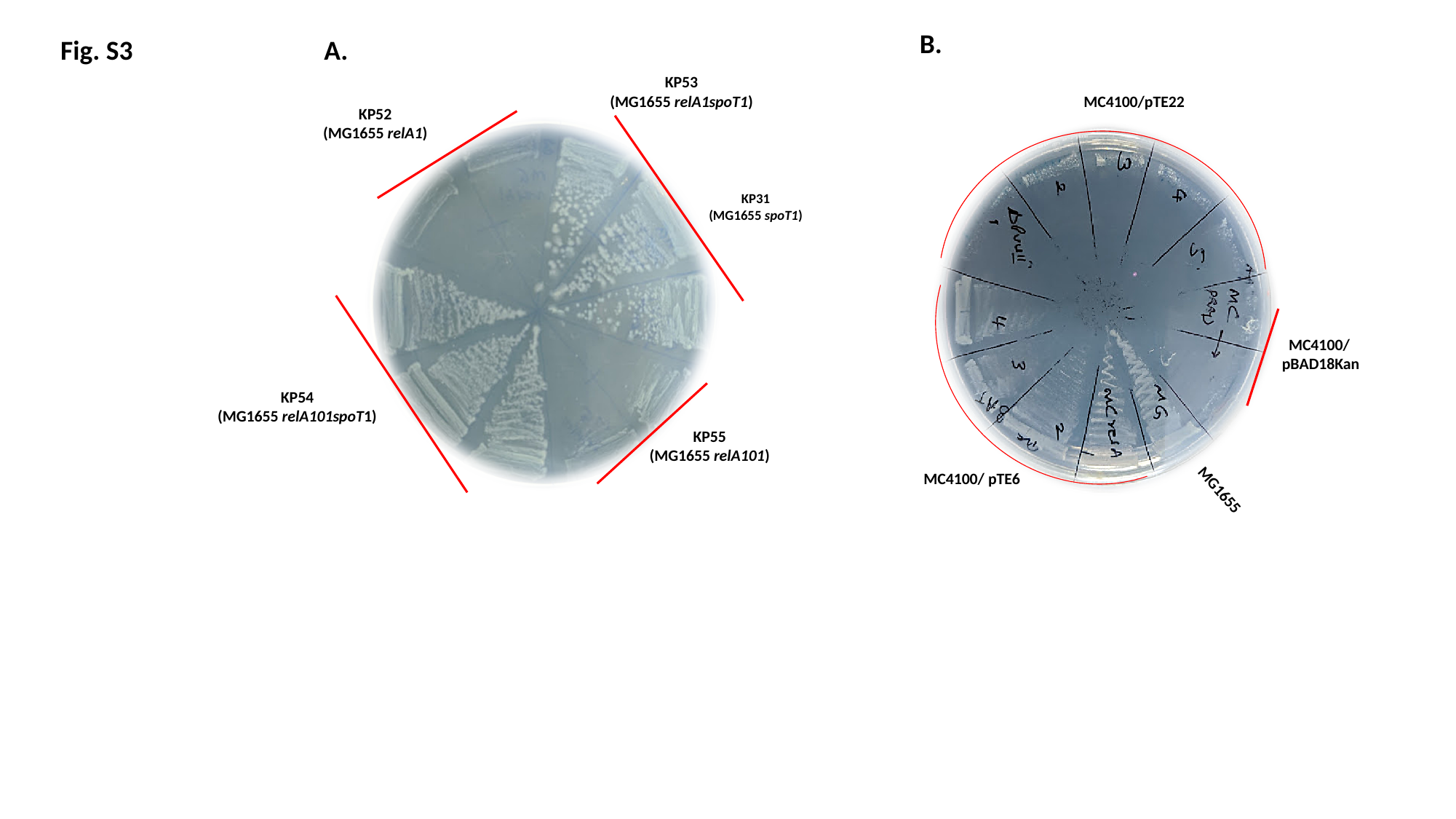

B.
A.
KP53
(MG1655 relA1spoT1)
MC4100/pTE22
KP52
(MG1655 relA1)
KP31
(MG1655 spoT1)
MC4100/
pBAD18Kan
KP54
(MG1655 relA101spoT1)
KP55
(MG1655 relA101)
MC4100/ pTE6
MG1655
Fig. S3

### Slide 3
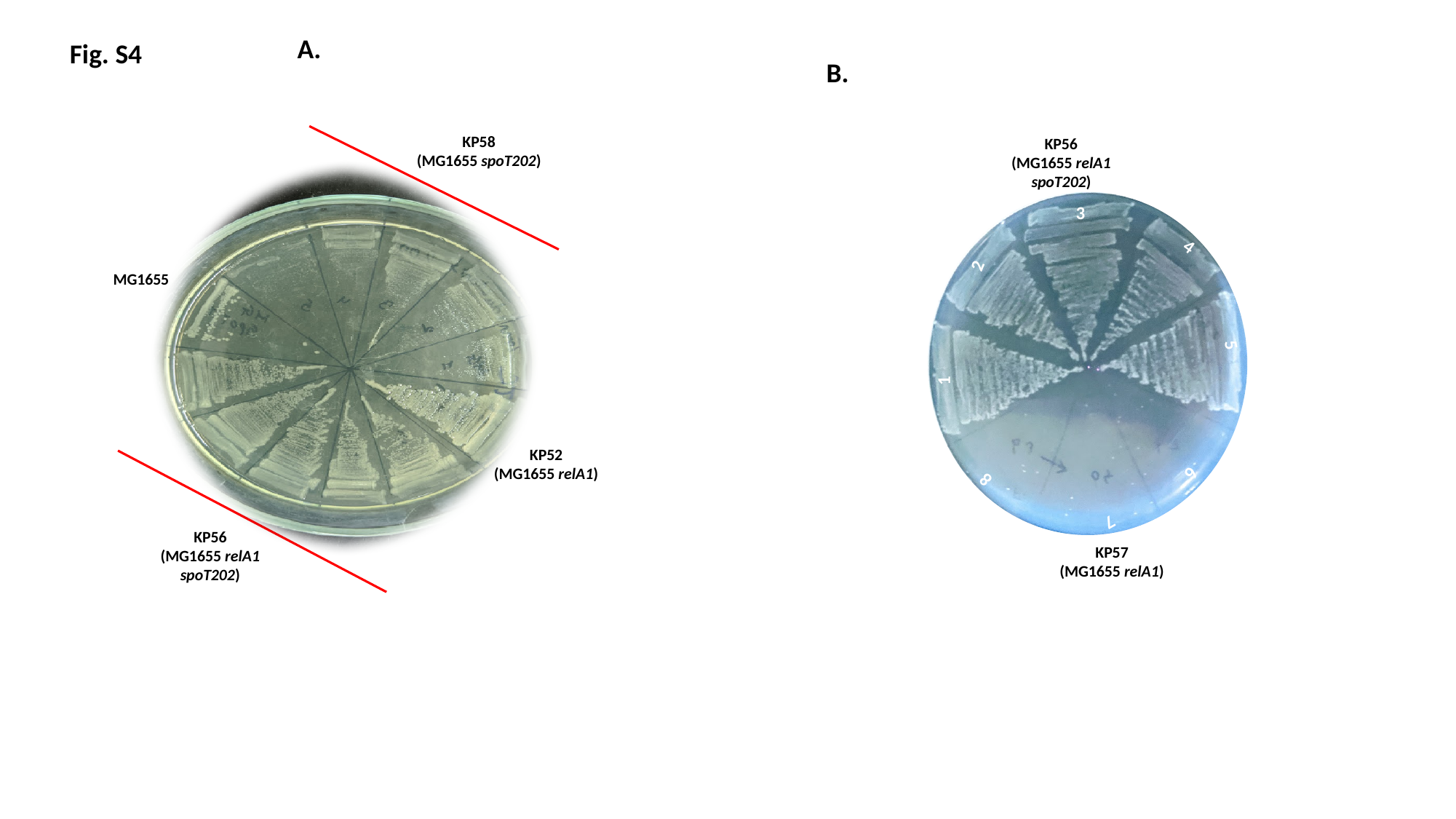

A.
Fig. S4
B.
KP58
(MG1655 spoT202)
KP56
(MG1655 relA1 spoT202)
3
4
2
MG1655
5
1
KP52
(MG1655 relA1)
6
8
7
KP56
(MG1655 relA1 spoT202)
KP57
(MG1655 relA1)
